# Sequence-based modeling of plant epigenomes reveals cell-type-specific *cis*-regulatory grammar

**DOI:** 10.64898/2026.07.22.740070

**Authors:** Jie Yao, Jiaqi Li, Xuan Zhang, Xiang Li, Alexandre P. Marand, Ethan Pickering, Robert J. Schmitz

## Abstract

How *cis*-regulatory sequences and their genetic variation govern chromatin accessibility and gene expression to establish plant cell identities remains incompletely understood, despite their fundamental roles in development, environmental responses, and phenotypic diversity. Here we present PEAgent, a framework for training, evaluating and interpreting deep-learning models that predict single-cell chromatin accessibility directly from DNA sequence, packaged in an interactive web portal and toolkit. Models were trained on single-cell chromatin-accessibility atlases of soybean, maize and rice, together spanning over 355,000 cells and 320 cell types and ∼150 million years of evolution. We unraveled a lexicon of 243 cell-type-resolved regulatory patterns, half of them composite, with TCP and bHLH showing the greatest influence and strongest conservation across species. Co-occurrence and *in silico* synergy analyses, explicitly modeling motif orientation and spacing, revealed two distinct cooperative modes acting at short and nucleosome-scale distances. We further showed that model predictions distinguish grass-conserved from rice-specific regulatory sequences far more accurately than sequence conservation scores alone, and validated the model’s predicted variant effects against cell-type-level chromatin accessible QTLs. PEAgent provides a foundational resource for decoding cell-type-specific plant *cis*-regulatory logic and interpreting noncoding variation in plants.

**Highlights:**

- PEAgent predicts single-cell chromatin accessibility directly from DNA sequence in soybean, rice and maize.
- Nearly half of cell-type-resolved patterns are composite, with TCP and bHLH motifs most influential and conserved across species.
- Co-occurrence and synergy analysis reveals distinct cooperative grammar at short and nucleosome-scale distances.
- PEAgent distinguishes grass-conserved from lineage-specific regulatory sequences better than sequence conservation scores alone.

## Introduction

Plant phenotypic diversity is shaped not only by protein-coding variation but also by sequence variation in *cis*-regulatory elements that determine the timing, spatial pattern, and magnitude of gene expression^1–3^. These sequence elements are typically found within accessible chromatin regions (ACRs), where nucleosome depletion is often associated with transcription-factor binding^4^. *Cis*-regulatory grammar refers to the number, spacing, orientation and arrangement of transcription-factor binding sites within a regulatory sequence, and to how this organization influences chromatin accessibility and gene expression^5^. *Cis*-regulatory sequence conservation and divergence are central to phenotypic evolution^6,7^. Recent comparative genomics analysis across broad plant diversity has revealed widespread deep conservation alongside lineage-specific turnover of noncoding regulatory sequences^8^. However, sequence conservation alone does not explain how these sequence elements encode cell-type-specific regulatory activity or how individual genetic variants reshape that activity. Although the genome is shared across the cells of an individual, regulatory activity is deployed in a cell-type-specific manner, generating distinct cell states and identities^8^. Understanding how noncoding sequences encode chromatin accessibility, and how variants perturb this process within specific cell types, is therefore central to interpreting agronomically important regulatory variation and its evolution. Yet, the sequence grammar governing cell-type-resolved chromatin accessibility in plants remains poorly understood, limiting our ability to connect noncoding variation to regulatory function and phenotype.

Deep-learning models can now predict functional genomic signals directly from DNA sequence, an approach termed sequence-to-function modeling that learns the regulatory code from data rather than from predefined motifs^10,11^. This paradigm has advanced fastest in mammals, where large, high-quality single-cell datasets are available. Models such as scBasset^12^ and chromBPNet^13^ predict chromatin accessibility from sequence at single-cell and cell-type resolution in animals, and such predictions have been extended to reconstruct cell-type-resolved regulatory landscapes across vertebrate species, to infer *cis*-regulatory evolution (DeepCREvo)^14^, and to interpret noncoding disease variants in specific human cell types^15^. In plants, deep-learning approaches have also been applied to predict molecular or phenotypic traits from DNA sequence^16–18^, but these efforts have largely remained at the bulk level, averaging signals across heterogeneous cell types and obscuring the cell-type-specific regulatory grammar that shapes development and trait variation. This gap is rooted in data: plants have lacked the large-scale, high-quality training datasets available for human and animal systems, which has limited the dissection of cell-type-resolved regulatory grammar.

Here, we present PEAgent (Plant Epigenome Agent), a framework for training, evaluating and interpreting deep-learning models that predict single-cell chromatin accessibility directly from DNA sequence. We trained one model per species on single-cell assay for transposase-accessible chromatin sequencing (scATAC-seq) atlases of soybean, maize and rice, together spanning more than 355,000 nuclei and 320 cell types. Interpreting these models, we resolved a lexicon of 243 cell-type-resolved regulatory patterns, nearly half of them composite motifs, with TCP and bHLH family motifs the most influential and cross-species-conserved determinants of chromatin accessibility. Co-occurrence and *in silico* synergy analyses uncovered two cell-type-specific cooperative modes acting at short and nucleosome-scale distances. The models also distinguished rice-lineage-specific from conserved ACRs in grasses better than sequence-conservation scores, and their predicted variant effects were validated against cell-type-level chromatin accessible QTLs (caQTLs). Finally, we provide these models, interpretations and tools through an interactive web portal (https://peagent.org/) for decoding *cis-* regulatory logic and prioritizing functional noncoding variation in plants.

## Results

### Construction and evaluation of a sequence-based model for plant single-cell chromatin accessibility

To decode *cis*-regulatory sequence features predictive of chromatin accessibility across heterogeneous plant cells, we used a deep-learning strategy^12,19^ to build predictive models from DNA sequence to single-cell chromatin accessibility in plants. We adapted the sequence-based convolutional neural network scBasset^12^ and built it into PEAgent (Plant Epigenome Agent), a framework for training, evaluating and interpreting these models (**Figure 1A**; **Methods**). The model takes a one-hot-encoded 1,344-bp sequence centered on an ACR as input and learns from a binarized cell × ACR accessibility matrix in a multi-task classification setting to predict a per-cell probability of chromatin accessibility in [0,1]. We trained an independent model on the scATAC-seq atlas of each of three species spanning diverse genomic architectures and evolutionary histories: maize (*Zea mays*; 50,632 cells; 158,586 ACRs, 94 cell types)^20^, rice (*Oryza sativa*; 104,029 cells; 128,764 ACRs, 128 cell types)^21^ and soybean (*Glycine max*; 200,732 cells; 303,199 ACRs, 98 cell types)^22^.

**Figure 1.**
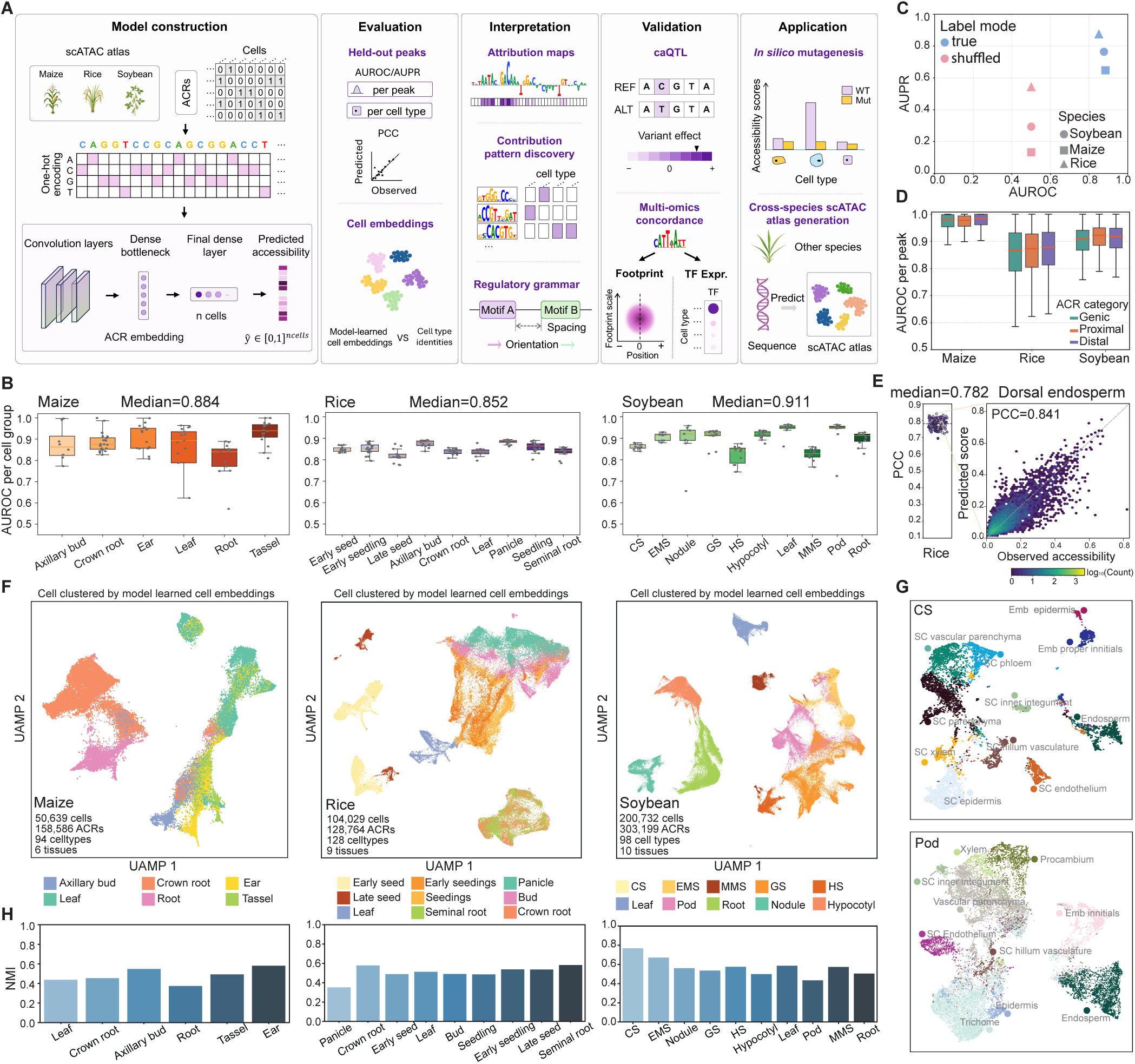
Sequence-based modelling of plant single-cell chromatin accessibility. **A**, Overview of the PEAgent framework for predicting single-cell chromatin accessibility from DNA sequence, comprising model construction, evaluation, interpretation, validation and application. **B**, Per-cell-group AUROC on held-out ACRs in soybean, maize, and rice. Each point represents one cell group in this tissue. Soybean tissues include GS, globular stage; HS, heart stage; CS, cotyledon stage; EMS, early maturation stage; and MMS, middle maturation stage. **C,** Comparison of performance using true or peak-shuffled labels, shown by median AUROC and AUPR for each species. **D,** Per-peak AUROC stratified by ACR category. Proximal and distal ACRs were defined relative to the nearest annotated gene using a 2-kb threshold. **E,** Pearson correlation coefficient (PCC) between predicted and observed chromatin accessibility on held-out peaks across cell type in rice, right panel show instance from dorsal endosperm. **F,** UMAP visualization of model-learned cell embeddings in soybean, maize and rice, colored by tissue. **G,** Higher-resolution UMAP views of cotyledon-stage seed (CS) and pod tissues in soybean. colored by cell type. **H**, Normalized mutual information (NMI) between embedding-derived Leiden clusters and annotated cell identities across tissues. NMI = 0 indicates random agreement.

We evaluated model performance from four perspectives: held-out ACR prediction accuracy, correlation between predicted and observed cell-type chromatin accessibility profiles, performance across different ACR classes, and the biological information encoded in the model-learned cell embeddings (**Methods**). On held-out ACRs unseen during training, PEAgent accurately predicted cell type level chromatin accessibility, measured by the area under the receiver operating characteristic curve (AUROC), across all three species (median AUROC per cell group: maize = 0.884, rice = 0.852, soybean = 0.911; **Figure 1B and TableS1**). At the individual peak level, per held-out peak AUROC reached 0.959, 0.898, and 0.812 in maize, soybean, and rice, respectively (**Figure S1A**). Both AUROC and the area under the precision-recall curve (AUPR) performance collapsed upon label shuffling (**Figure 1C**), confirming that predictions reflect sequence-encoded determinants of chromatin accessibility rather than dataset-level confounders.

We examined whether predictive accuracy varied across ACR categories. Performance remained broadly comparable across genic, proximal (≤2 kb), and distal (>2 kb) ACR categories in all three species (**Figures 1D and S1B**), indicating that PEAgent captures predictive sequence features irrespective of proximity to annotated genes. When ACRs were instead stratified by breadth of chromatin accessibility, both constitutively accessible ACRs (present in >80% of cell groups) and cell-type-specific ACRs (ctACRs) were predicted accurately, with modestly higher accuracy for constitutive ACRs in soybean and rice (**Figure S1C, Methods**). PEAgent also quantitatively recapitulated the magnitude of chromatin accessibility, with median Pearson correlation coefficient (PCC) on held-out peaks between predicted score and observed chromatin accessibility reaching 0.730, 0.782, and 0.761 in maize, soybean, and rice (**Figures 1E, S1D and S1E**).

We further asked whether the models learned biologically meaningful representations of cell identity. The final layer of PEAgent assigns each profiled cell a low-dimensional embedding, learned solely from predicting chromatin accessibility from sequence, given no explicit information about relationships among cell types. UMAP visualization of model-learned cell embeddings revealed clear tissue- and cell-type-level organization across all three species (**Figures 1F, 1G, and S1F**). To quantify this organization, we grouped cells by their model-learned embeddings using Leiden clustering and measured how closely these sequence-derived clusters matched the annotated cell identities with normalized mutual information (NMI) (**Methods**). NMI confirmed strong concordance across all tissues examined (**Figure 1H**). In soybean, NMI ranged from 0.768 (cotyledon-stage seed) to 0.434 (pod), yet even the lowest-NMI tissue exhibited clear cell-type stratification in UMAP space (**Figure 1G**). Together, these results demonstrate that PEAgent learns sequence representations that capture the biological organization of plant cell types.

### *In silico* mutagenesis reveals plant *cis*-regulatory sequence features associated with chromatin accessibility

To identify the *cis*-regulatory sequence features that influence chromatin accessibility, we interrogated the trained PEAgent models by *in silico* mutagenesis (ISM). For each ACR, we substituted every position with all alternative nucleotides and recorded the effect on the model’s bottleneck representation (**Methods**). Propagating these effects to the output layer produced reference-sequence attribution maps at two scales: a global attribution map averaged across all cells, and cell-type-specific attribution maps (**Figure 2A**). From these maps, TF-MoDISco^22^ extracted seqlets (short subsequences of high attribution) and clustered recurrent seqlets into contribution patterns (*de novo* motifs predictive of chromatin accessibility), each summarized as a contribution weight matrix and expected to correspond to a transcription-factor binding motif^10,12,15^. This recovered global patterns from constitutively accessible ACRs and cell-type-resolved patterns from ctACRs.

**Figure 2.**
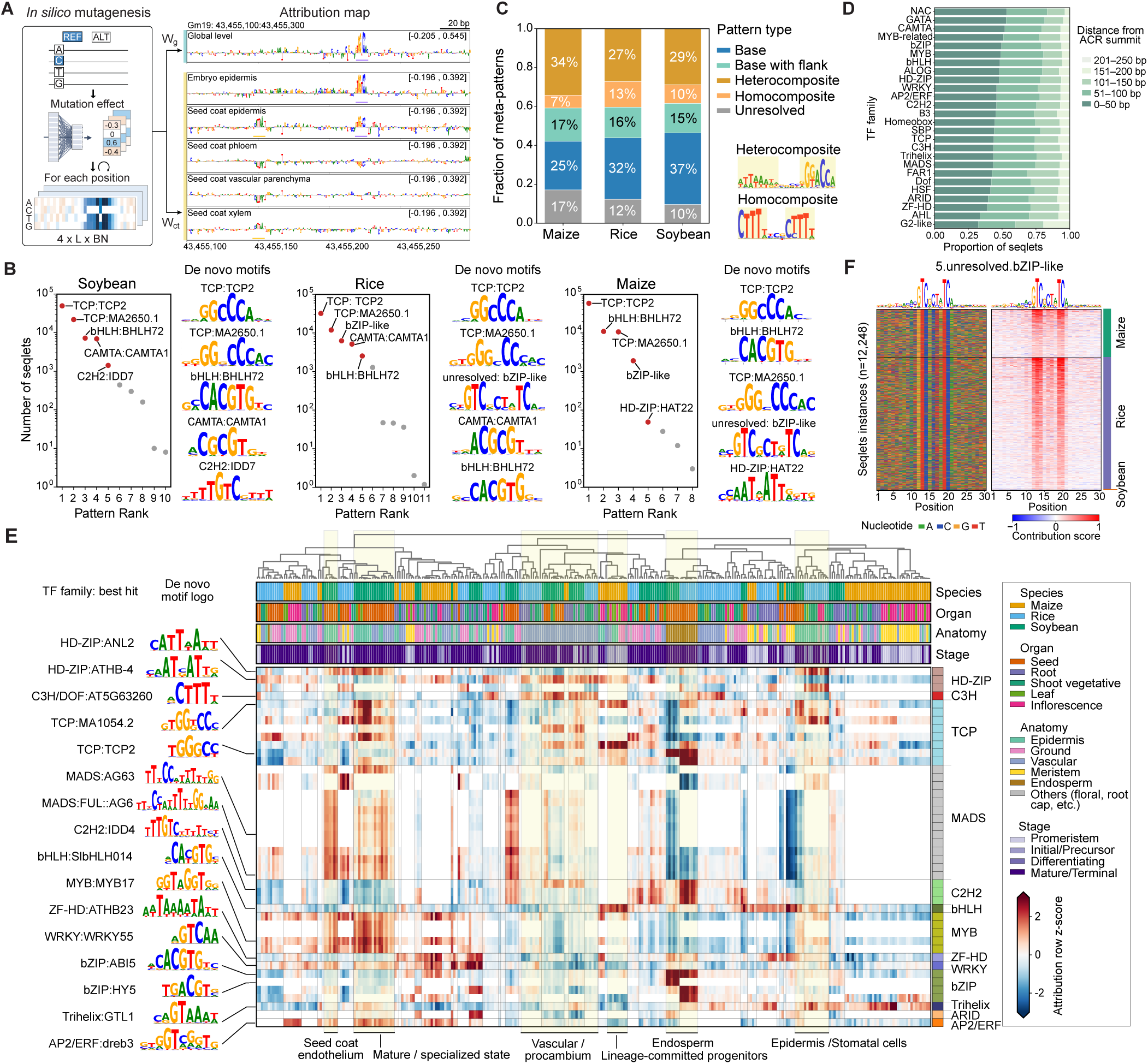
Model interpretation reveals shared and cell-type-specific *cis*-regulatory sequence contribution patterns across plants. **A**, Schematic of the ISM-based attribution analysis pipeline. Each nucleotide in a 1,344-bp ACR-centred detection window was substituted with alternative bases, and bottleneck effects were projected onto global (bulk-level, averaged across all cells) or cell-type-specific output weights. Attribution scores indicate the contribution of the reference nucleotide to predicted chromatin accessibility. Representative tracks are shown at Gm19:43,455,100-43,455,300. **B,** Global contribution patterns recovered from constitutively accessible ACRs in each species. Points rank TF-MoDISco-identified patterns by the number of seqlets; labels indicate TF family:best-hit motif annotations or unresolved patterns (q > 0.1). Logos show the top five contribution-weight matrices (CWMs) per species. **C,** Classification of 243 cell-type-resolved meta-patterns into base, base with flank, heterocomposite, homocomposite and unresolved classes, with representative composite logos. **D,** Distances of seqlets from cell-type-resolved base meta-patterns to ACR summits, summarized by TF family. **E,** Attribution landscape of all base meta-patterns across plant cell types. Rows show TF-family-grouped meta-patterns with representative CWM logos and TF family:best-hit annotations; columns show cell types annotated by species, organ, anatomy and developmental stage. Heatmap values are row-scaled attribution z-scores. **F,** Nucleotide composition and contribution scores for an unresolved pattern with consensus GTCNCTATC. Seqlets (n = 12,248) are aligned by motif position and stratified by species.

We first focused on the global patterns, which summarize the sequence features predictive of chromatin accessibility across all cells. We recovered 10 global patterns in soybean, 11 in rice and 8 in maize in total. The patterns with the most supporting seqlets were highly conserved across species (**Figure 2B**). In each species, the most abundant pattern matched TCP-family motifs, along with the bHLH G-box (CACGTG; best match BHLH72). This is consistent with a previous population-scale maize caQTL study in which variants within TCP-binding sites were the most prevalent determinants of chromatin accessibility^24^. The recurrence of these dominant global patterns across the monocots maize and rice and the dicot soybean suggests a partially conserved *cis*-regulatory grammar underlying chromatin accessibility in flowering plants.

Next, we characterized cell-type-resolved motifs. We pooled the contribution patterns initially recovered across all cell types and species, then clustered, denoised and classified them (**Methods**). This yielded 243 meta-patterns spanning 262 cell types across soybean, rice and maize. This set recovered the globally defined contribution patterns and substantially expanded on them with cell-type-restricted patterns. Guided by the motif-classification framework of Liu et al.^15^, we assigned these meta-patterns into five classes (**Figure 2C and Table S2, Methods**). Base motifs (n = 57) matched known transcription-factor binding motifs in databases, and base-with-flanks motifs (n = 47) extended these with additional predictive flanking nucleotides. Base motifs were the most evolutionarily constrained class, with 53% shared across multiple species, the highest proportion of any class (**Figures S2A and S2B**). Notably, over 47% of meta-patterns were composite motifs, each comprising two sites of the same (homocomposites) or different (heterocomposites) TF family, and showed higher cell-type specificity than base motifs (**Figures 2C and S2C**). The remaining were classified as ‘unresolved’ (n = 30) because they could not be confidently matched to any known TF-binding motif (q > 0.1). We next examined the genomic location of these cell-type-resolved meta-patterns. Across soybean, maize and rice, the seqlet instances were enriched toward ACR centers, mirroring the density distribution of the global patterns (**Figures 2D and S2D**). Among the TF families represented by base motifs, the TCP and AP2/ERF families showed remarkably broad and more distal distributions relative to the TSS (TCP median 6.2 kb, Q3 24.6 kb; AP2/ERF median 5.4 kb), whereas the seqlets of the remaining families had median TSS distances of 2.4 to 3.1 kb (**Figure S2E**). This distal bias is consistent with previous reports that TCP-binding sites are enriched at gene-distal enhancers and long-range chromatin interactions in rice and maize^20,24,25^.

Meta-patterns matching known transcription factors showed divergent cell-type specificity in their attribution scores across TF families, in line with the known enrichment of these motifs in ctACRs (**Figure 2E**). For example, the TCP motif (TGGCC; best hit TCP2) and the bZIP G-box (best hit ABI5) showed their strongest attribution signals in endosperm across species. These distributions recover what single-cell experiments report directly, with TCP and bZIP motifs enriched in endosperm-specific unmethylated ACRs in rice^21^, and the ABI5 motif showing endosperm-biased deviation in soybean scATAC profiling^22^. The C2H2 meta-pattern (best hit IDD4) was similarly endosperm-restricted, in agreement with its enrichment in dorsal and lateral starchy endosperm in the rice atlas^21^. HD-ZIP motifs (best hit ANL2) showed their highest attribution in epidermal and vascular cells at the mature developmental stage, matching both their cell-type distribution in the soybean^22^ and the established role of ANL2-type HD-ZIP IV factors in patterning the epidermal and subepidermal layers^26^. The C3H/DOF meta-pattern (CTTTT) displayed its highest attribution in procambial and other vasculature-related cells across species, paralleling the vascular-specific enrichment of DOF motifs in soybean single-cell data^22^. Together, this concordance between deep-learning attribution and independent motif-enrichment analysis indicates that PEAgent captured the cell-type *cis*-regulatory grammar and assigned it to the correct cellular context.

Not every predictive contribution pattern corresponded to a known transcription factor. The most common unresolved pattern (consensus GTCNCTNTC) was also recovered among the global patterns and was especially abundant in maize and rice (**Figure 2F**). It carried no distinguishing genomic features, with a distribution indistinguishable from ACRs overall and from other annotated TF motifs and no enrichment in gene features (UTRs, introns and so on) or transposable elements (**Figures S2F-S2H**). Nonetheless, footprint analysis revealed a clear signal centered on the motif, suggesting it is a binding site of a transcription factor (**Figure S2I**).

### Co-occurrence and synergy of contribution patterns reveal combinatorial *cis*-regulatory syntax

Combinatorial arrangements of transcription-factor binding sites stabilize binding through cooperative recruitment^27^, and by helping transcription factors outcompete nucleosomes, establish and maintain accessible chromatin^28^. Co-occurrence of contribution patterns within a single ACR was pervasive in our data, with 56.4% of ctACRs containing more than one seqlets (**Figure S3A**). To test whether these co-occurrences are spatially organized and cell-type-specific, we systematically tabulated the natural co-occurrence of all base-motif pairs across 1,596 pairwise combinations and 261,366 co-occurrence instances. The center-to-center distance between paired motifs was bimodal, with peaks at approximately 8 bp and 43 bp (**Figure 3A**; center-to-center and edge-to-edge distance metrics are defined in **Figure S3B**). These two spacing regimes may reflect two distinct modes of combinatorial action: a short-range mode (Type 1, ≤20 bp), in which adjacent transcription factors stabilize each other’s binding through direct contact or DNA-mediated allostery and cooperatively displace the local nucleosome, and a nucleosome-range mode (Type 2, 20–150 bp), still within a single nucleosome and consistent with collective transcription-factor competition against the nucleosome^15,29^.

**Figure 3.**
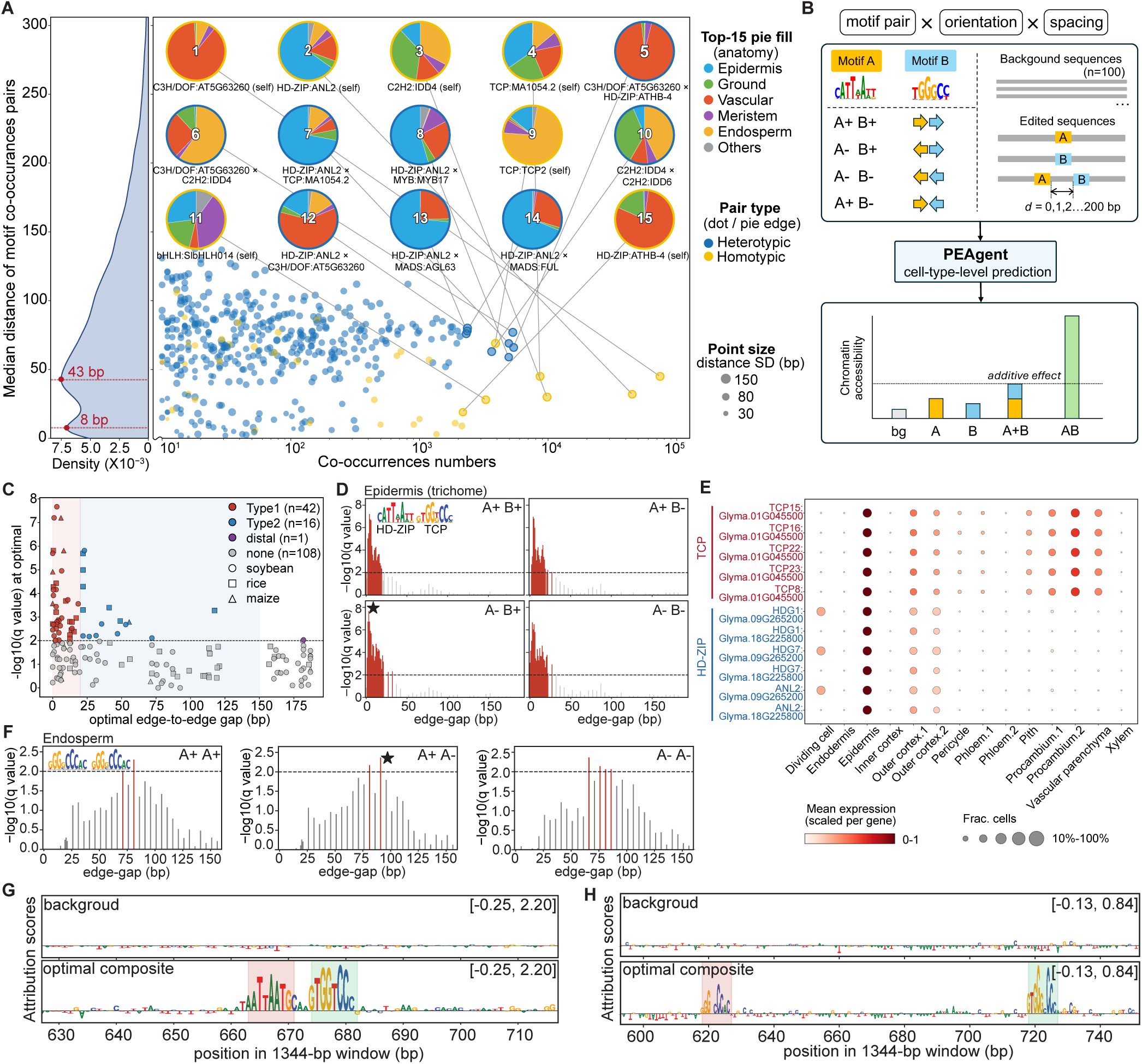
Co-occurrence and synergy of contribution patterns. **A,** Co-occurrence of base-motif pairs (one point per pair). x axis, number of co-occurrence instances; y axis, median center-to-center distance; color, pair type (heterotypic, blue; homotypic, yellow); point size, distance s.d. Left, density of center-to-center distances over all co-occurrence instances. Pies, the 15 most frequent pairs; fill, anatomical distribution of instances; edge, pair type. **B,** Schematic of the *in-silico* marginalization used to quantify synergy. For each motif pair, motif A and motif B are inserted alone and together into 100 background sequences across all four relative orientations and spacings (d = 0 to 200 bp). **C,** Synergy of all tested pairs. x axis, optimal edge-to-edge gap; y axis, −log10 q at the optimal arrangement. q, Benjamini–Hochberg-adjusted P value. Dashed line, q = 0.01. **D,** Synergy of the HD-ZIP and TCP pair in the epidermis across the four relative orientations and edge-gaps. star, optimal arrangement. **E,** Single-cell RNA-seq expression of TCP and HD-ZIP family members across cell types of the soybean hypocotyl. **F,** Synergy of a homotypic TCP pair in the endosperm across orientations and edge-gaps. star, optimal arrangement. **G,** Sequence attribution for the optimal HD-ZIP and TCP composite (bottom) compared with background (top). **H,** Sequence attribution for the optimal endosperm TCP composite (bottom) compared with background (top).

The most frequent co-occurring pairs comprised both homotypic (two copies of the same motif; 7 of the top 15) and heterotypic (two different motifs; 8 of the top 15) combinations, most of which showed a pronounced cell-lineage preference (**Figure 3A**). For example, homotypic DOF pairs (no.1) and heterotypic HD-ZIP × DOF pairs (no.5) occurred predominantly in vascular tissue, whereas heterotypic HD-ZIP–TCP pairs (no. 7) were enriched in the epidermis. This lineage preference was corroborated by independent single-cell RNA-seq data, in which the transcription factors underlying each pair were co-expressed in their corresponding lineages (**Figures 3E and S3C**).

To distinguish genuine synergy from simple co-occurrence, we asked whether motif pairs act synergistically, with a joint effect on chromatin accessibility exceeding the sum of their individual effects, and whether such synergy is constrained by orientation and spacing. Using PEAgent models, we quantified synergy between motif pairs by *in silico* marginalization strategy at cell-type resolution^15^. For each pair, we inserted the two motifs separately (yielding single-motif effects A and B) and together (yielding the joint effect AB) into 100 background sequences of low baseline chromatin accessibility and controlled GC content. A pair was scored as synergistic when its joint effect (AB) significantly exceeded the additive expectation (A + B), tested across all four relative orientations and edge-to-edge (e2e) gaps of 0 to 200 bp (**Figure 3B**, **Methods**). Testing each pair in the cell type where it was most abundant, restricted to pairs with more than 20 instances, we identified 59 synergistic pairs among the 167 examined (q < 0.01; **Figure 3C and Table S3**). Most synergy was short-range, with 42 pairs having optimal e2e gaps below 20 bp (Type 1) and the remaining at longer distance, mostly within a single nucleosome scale (Type 2). Overall, 71% of synergistic pairs were sensitive to the relative orientation of the two motifs (**Figure S3D**), indicating that a defined angular arrangement, and not proximity alone, is often required for greater-than-additive activity.

Several synergistic pairs recapitulated transcription-factor interactions with independent experimental support. For example, an HD-ZIP and TCP pair showed the strongest synergy in the epidermis at a short optimal e2e gap of about 3 bp and was insensitive to relative orientation (**Figures 3D, 3G and S3E**). Consistent with a shared epidermal role, HD-ZIP factors, including ANL2 (the best hit for this HD-ZIP motif), and several TCP factors were specifically co-expressed in the soybean epidermis in single-cell RNA-seq data (**Figure 3E**), and ANL2 is a known regulator of epidermal cell differentiation^26,30^. The predicted synergy is further supported by direct evidence in Arabidopsis, where TCP and HD-ZIP factors cooperatively activate an epidermal enhancer, as shown by enhancer dissection and reporter analysis^31^. Mechanistically, yeast two-hybrid and bimolecular fluorescence complementation assays show that HD-ZIP proteins physically interact with CIN-type TCP factors through their C-terminal leucine zipper^32^. In a parallel, endosperm-focused analysis of all 36 co-occurring pairs, we identified a synergistic homotypic TCP pair. Rather than acting at short range, this pair reached maximal synergy at a broad spacing of about 65 to 90 bp (**Figures 3F, 3H and Table S4**), a distance too large for direct protein-protein contact but well within a single nucleosome^33^, pointing to a nucleosome-scale mode (Type 2). This long-range mode is consistent with the known ability of TCP factors to recruit the SWI/SNF chromatin-remodelling ATPase BRAHMA to their targets, a nucleosome-scale activity that could allow two nearby TCP sites to influence the same nucleosome cooperatively rather than through direct contact^34^.

### Cross-species prediction of single-cell chromatin accessibility and evolutionary classification of rice *cis*-regulatory sequences

Generating plant scATAC-seq atlases remains challenging due to costs, and because some species and tissues are technically difficult to process, limiting cell-type-resolved studies of chromatin accessibility and *cis*-regulatory elements. By predicting chromatin accessibility directly from DNA sequence, PEAgent can reconstruct chromatin accessibility landscapes for species that lack their own scATAC atlases, potentially circumventing these barriers. To assess how well the models transfer across species, we evaluated all nine within- and cross-species prediction combinations among the independently trained maize, rice, and soybean models. PEAgent showed robust cross-species transferability, with bulk-level AUROC values of 0.725–0.880 and cell-type-level median AUROC values of 0.713–0.853 across all cross-species pairs **(Figure 4A)**. Prediction accuracy was highest between the closely related monocots rice and maize, whereas transfers involving soybean showed lower performance, consistent with greater evolutionary divergence.

**Figure 4.**
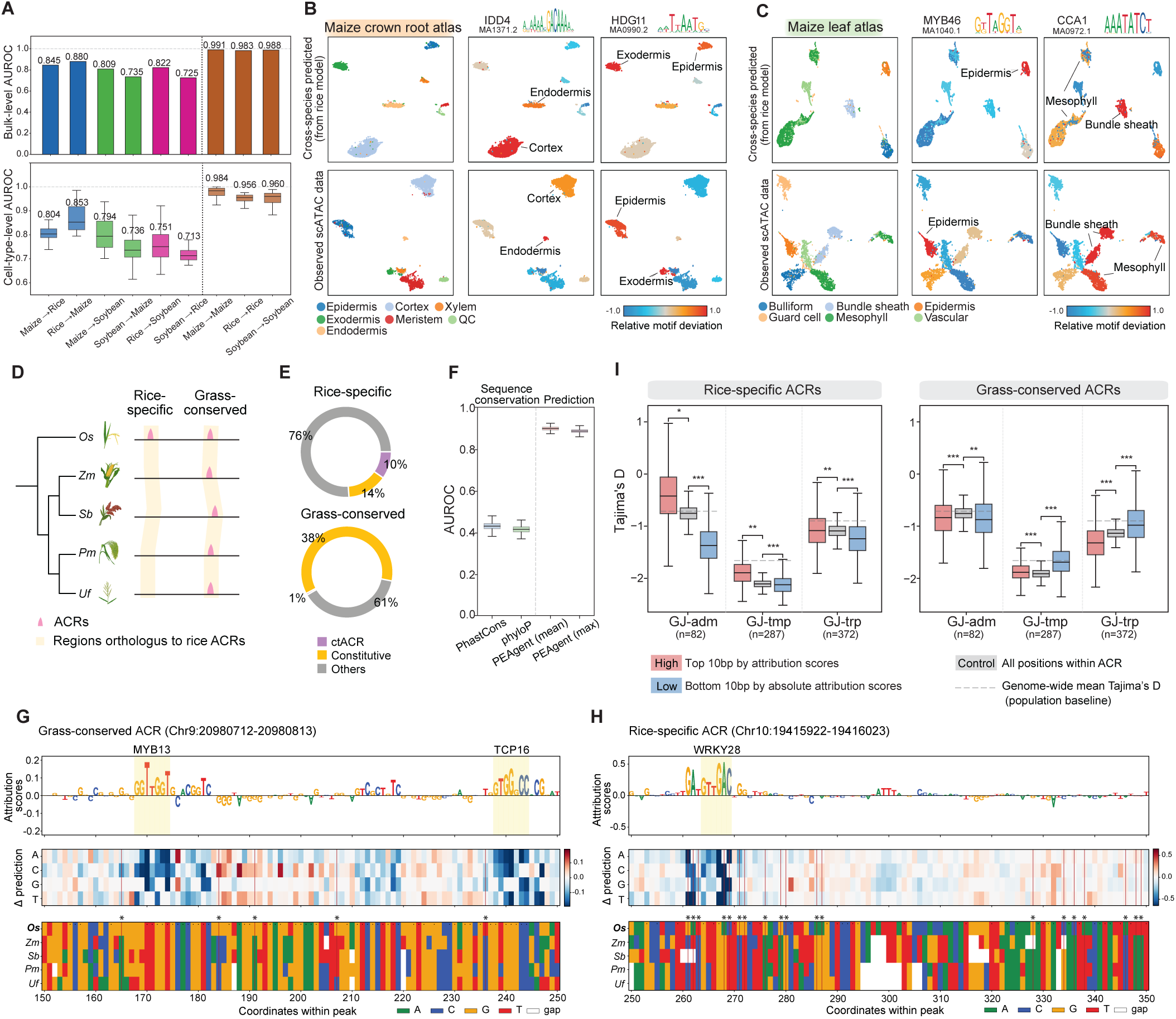
Cross-species prediction of chromatin accessibility and evolutionary stratification of rice *cis*-regulatory elements. **A**, Bulk-level (top) and cell-type-level (bottom) AUROC for all nine within- and cross-species prediction pairs among maize, soybean, and rice. **B-C,** Comparison of predicted and observed maize crown root (**B**) and leaf (**C**) scATAC-seq atlases. Top panels show rice-trained model predictions; bottom panels show observed maize data. Representative TF motif deviation scores are averaged by cell type. **D,** A phylogenetic tree of five grass species illustrates the classification of rice ACRs as rice-specific or grass-conserved based on chromatin accessibility within orthologous sequence regions. *Os*, *Oryza sativa*; *Zm*, *Zea mays*; *Sb*, *Sorghum bicolor*; *Pm*, *Panicum miliaceum*; *Uf*, *Urochloa fusca*. **E,** Proportions of cell-type-specific (ctACR), constitutively accessible, and other ACRs within rice-specific and grass-conserved categories. **F,** Classification performance (AUROC, bootstrap, 1,000 iterations) of sequence conservation scores (PhastCons, PhyloP) versus PEAgent predictions averaged across orthologues grass sequences (cell-type mean and maximum in leaf) in distinguishing rice-specific and grass-conserved ACRs. **G-H,** Instance of grass-conserved **(G)** and rice-specific **(H)** ACRs illustrating attribution score profiles (top), *in silico* mutagenesis variant-effect heatmaps (middle), and multispecies sequence alignments across *Os*, *Zm*, *Sb*, *Pm*, and *Uf* (bottom). **I,** Distributions of Tajima’s D (bootstrap, 1,000 iterations across ACRs) for positions with high (top 10 bp by attribution score), low (bottom 10 bp by absolute attribution score), and control (all positions within ACR) model contribution within rice-specific and grass-conserved ACRs, across three japonica rice subpopulations(GJ-tmp, temperate; GJ-trp, tropical; GJ-adm, admixed). Dashed line indicates genome-wide mean Tajima’s D as a population baseline.

To further assess whether cross-species predictions preserved biologically meaningful cellular organization and regulatory specificity, we focused on rice-to-maize transfer, the best-performing cross-species setting. Predicted chromatin accessibility landscapes for maize crown root and leaf recapitulated the major cellular compartments of the observed atlases, with comparable UMAP organization across epidermal, ground tissue, and vascular cell groups (**Figures 4B and 4C**). Representative known transcription factor motifs also showed concordant cell-type-associated deviation patterns between predicted and observed atlases. For instance, the MYB46 motif, associated with secondary cell wall thickening in epidermal trichomes, was consistently enriched in epidermal cells in both predicted and observed maize leaf atlases (**Figures 4B, 4C, and S4A**). A genome browser screenshot illustrates concordant chromatin accessibility between cross-species prediction and maize observation at individual ACR loci across cell types (**Figure S4B**). These results show that PEAgent’s models exploit the conserved *cis*-regulatory grammar of plants^35^ to recover cell-type-specific chromatin features across species.

We next investigated whether incorporating PEAgent’s predictions can improve inference of ACR evolutionary histories compared with traditional sequence-based phylogenetic methods. Using syntenic region mappings and chromatin accessibility profiles across five grass species that we previously established (*Oryza sativa*, *Zea mays*, *Sorghum bicolor*, *Panicum miliaceum*, *Urochloa fusca*)^21^, we classified rice ACRs as grass-conserved or rice-specific based on the presence or absence of chromatin accessibility within syntenic regions, where grass-conserved ACRs were accessible across all these grass species whereas rice-specific ACRs were exclusively present in rice **(Figure 4D)**. Compared with rice-specific ACRs, grass-conserved ACRs were enriched for constitutively accessible ACRs (38% vs 14%), showing broader chromatin accessibility across cell types **(Figure 4E)**. Sequence conservation scores (PhyloP, PhastCons)^36,37^, which capture evolutionary constraint on primary sequence but cannot distinguish whether sequence divergence disrupts regulatory activity, served as baseline classifiers. PEAgent prediction scores, summarized as the mean or maximum chromatin accessibility across leaf cell types in grass syntenic sequences, substantially outperformed sequence conservation in classifying rice-specific versus grass-conserved ACRs (PEAgent mean AUROC = 0.897, PEAgent max = 0.885, versus PhastCons = 0.429, PhyloP = 0.414; **Figures 4F, S4C, S4D, and Table S5**). This indicates that sequence-to-accessibility deep-learning model predictions capture regulatory activity conservation beyond what is encoded in primary sequence similarity alone.

PEAgent’s capacity to capture the impact of precise sequence substitutions likely contributes to this superior classification: rather than tracking broad sequence divergence, the model distinguishes substitutions that disrupt critical regulatory features from those accumulating in neutral positions. To examine this mechanistically, we illustrated attribution maps and *in silico* mutagenesis variant-effect landscapes for representative ACRs of each class. For a grass-conserved ACR (Chr9:20980712– 20980813), rice-specific sequence changes fell outside high-attribution positions, which co-localized with MYB13 and TCP16 binding motifs **(Figure 4G)**. For a rice-specific ACR (Chr10:19415922– 19416023), multiple rice-specific substitutions instead fell at high-attribution positions best matching a WRKY28 motif **(Figure 4H)**. Consistent with these instances, rice-specific ACRs globally showed a significantly higher rice-specific change-associated positive attribution burden than grass-conserved ACRs (two-sided Mann-Whitney U test, BH-adjusted *P* = 8.7 × 10^−7^, **Figure S4E**). This mirrors observations in mammalian *cis*-regulatory evolution^14^, where the disruption of key functional positions by specific sequence changes, rather than broad sequence divergence, distinguishes lineage-specific from conserved regulatory elements.

To evaluate whether model attribution scores reflect positions with distinct evolutionary signatures, we examined Tajima’s D (a measure of the allele-frequency spectrum that is more negative under purifying selection) between high-attribution and low-attribution positions within rice-specific and grass-conserved ACRs, across three *japonica* rice populations^38^. Within grass-conserved ACRs, high-attribution positions showed similar or lower Tajima’s D than low-attribution positions across all three populations, consistent with stronger purifying selection at model-predicted functional sites. Within rice-specific ACRs, the pattern was reversed: high-attribution positions exhibited higher Tajima’s D than low-attribution positions, consistent with relaxed constraint at functionally important sites of recently emerged regulatory elements **(Figure 4I)**.

### PEAgent predicts regulatory variant effects validated by cell-type-resolved caQTL data

To evaluate PEAgent’s ability to predict the functional consequences of genetic variants on chromatin accessibility, we applied the model to 46,443 fine-mapped *cis*-caQTL single-nucleotide variants (SNVs) previously identified across 13 seedling cell types in 172 diverse maize inbred lines, using non-fine-mapped SNVs from the same ACR as non-causal controls. For each SNV, a sequence window of the model’s input length centered on the variant position was extracted, and the reference and alternate allele sequences were independently input to PEAgent. The variant effect was quantified as the difference in logit-transformed predicted chromatin accessibility between the two alleles (Δlogit, ALT minus REF, **Figure 5A**). Predicted variant effect magnitudes were significantly greater for fine-mapped candidate causal variants compared with same-ACR non-causal controls across 92.3% cell types (**Figure 5B**), suggesting that the model captures sequence features predictive of causal regulatory activity within accessible chromatin. We further found that directional accuracy was significantly higher for large-effect than small-effect variants in both the zero-shot maize model and B73 seedling fine-tuned maize models (top vs. bottom 20% by effect size; Wilcoxon test, p = 5.0 × 10⁻⁴ and p = 1.2 × 10⁻³, respectively; **Figure S5A**). Genome-wide visualization of alleles with large predicted variant effects revealed comparable cell-type-resolved patterns between the zero-shot and B73 seedling fine-tuned models, further supporting the reliability of PEAgent for direct genetic variant-effect prediction (**Figure S5B**).

**Figure 5.**
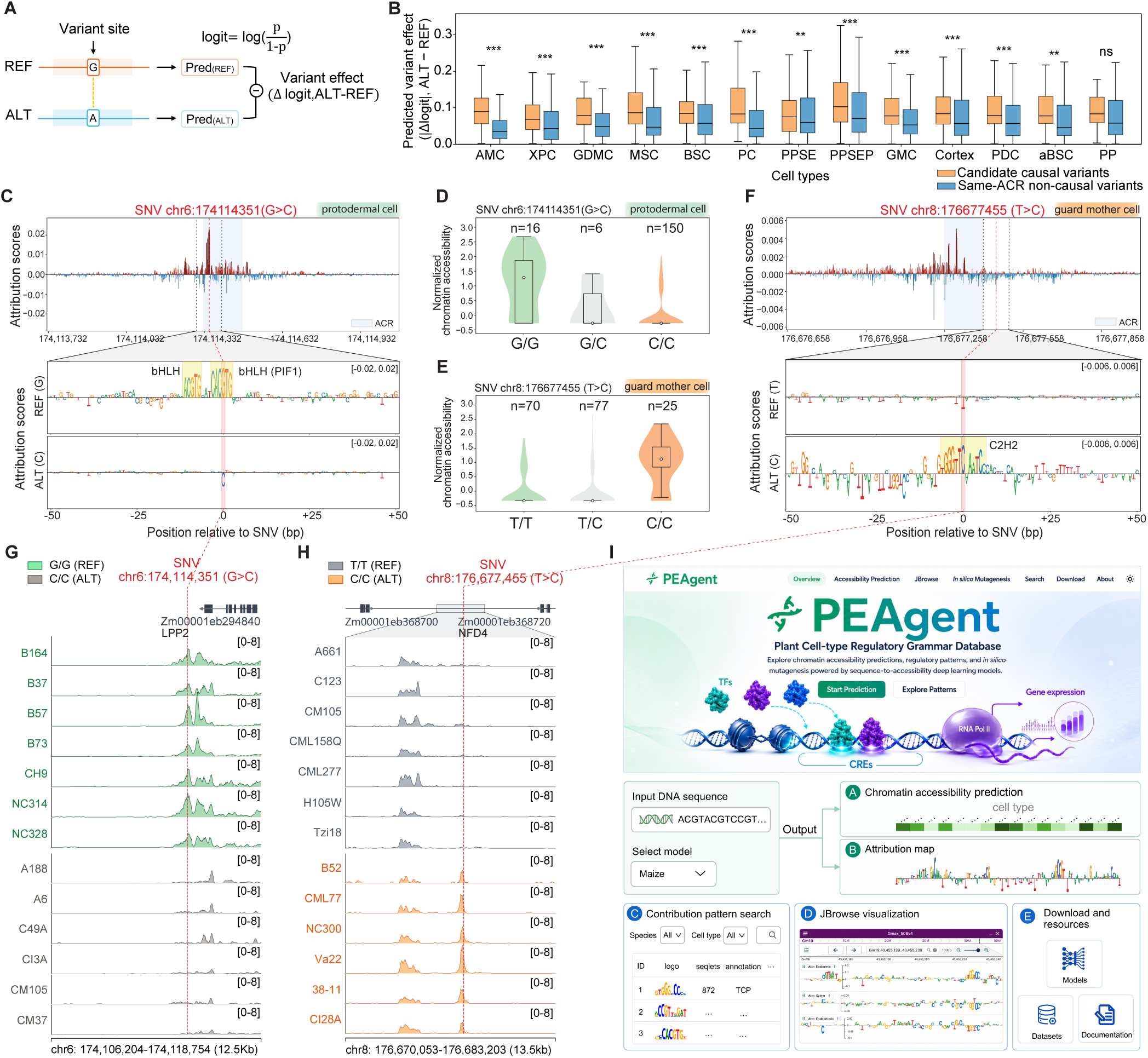
PEAgent predicts genetic variant effects on chromatin accessibility across cell types. **A**, Schematic of the center-mask variant effect scoring strategy for chromatin accessibility prediction. **B**, Predicted variant effect magnitudes for fine-mapped causal variants and four same-ACR non-causal controls across maize seedling cell types (Wilcoxon rank-sum test; ** p < 0.01, *** p < 0.001). **C**, Attribution score profile for a protodermal cell caQTL (SNV chr6:174,114,351, G>C). Top: attribution scores across the 1,344-bp *in silico* mutagenesis scanning window, with the population-defined ACR highlighted in blue. Bottom: per-nucleotide attribution scores for REF and ALT alleles within a ±50-bp window, showing the substitution disrupts a bHLH/PIF1 binding motif at a high-attribution position. **D-E**, Violin plots showing the distribution of normalized chromatin accessibility within the caQTL-associated ACR across three genotype classes at SNV chr6:174,114,351 (protodermal cells) and SNV chr8:176,677,455 (guard mother cells) in maize inbred lines. **F.** Attribution score profile for a guard mother cell caQTL (SNV chr8:176,677,455, T>C). Layout as in (c). The T>C substitution creates a C2H2 zinc finger motif at a high-attribution position, consistent with increased accessibility in the ALT allele. **G-H**, Genome browser tracks of chromatin accessibility at the chr6:174,114,351 and chr8:176,677,455 loci for 13 representative maize genotypes grouped by REF and ALT alleles. Tn5 integrations are scaled per million. **I**, Overview of the PEAgent web portal, showcasing its five core functionalities.

To illustrate the mechanistic basis of these predictions, we examined representative caQTLs in which PEAgent attribution scores provide interpretable links between sequence context and variant effect. For a protodermal cell caQTL (SNV chr6:174,114,351, G>C) in the promoter of the lipid phosphate phosphatase 2 gene (*LPP2*; Zm00001eb294840), attribution scores identified a bHLH/PIF1 binding motif as the dominant high-attribution element within the ACR; the G>C substitution fell precisely within this motif, abolishing it in the ALT allele (**Figure 5C**). Population-level scATAC-seq data confirmed significantly reduced chromatin accessibility in the caQTL-associated ACR in C/C genotypes within protodermal cells (**Figure 5D**), with genome browser tracks across individual inbred lines showing progressive peak attenuation from REF to ALT genotypes (**Figure 5G**). The guard mother cell caQTL (SNV chr8:176,677,455, T>C) illustrated the converse: the T>C substitution created a C2H2 zinc finger motif in the ALT allele (**Figure 5F**). Consistent with this motif gain, C/C genotypes showed elevated chromatin accessibility at the associated ACR in population data (**Figure 5E**), and genome browser tracks displayed progressively stronger peaks from REF to ALT genotypes (**Figure 5H**). These results demonstrate PEAgent’s capacity to predict both motif disruption and motif creation underlying regulatory variation. Two additional cortex caQTL examples show that SNVs disrupting only one of multiple high-attribution motifs within an ACR produce partial, graded reductions in chromatin accessibility rather than complete loss (**Figures S5C-S5H**).

To make PEAgent broadly accessible to the plant research community, we developed an interactive web portal (https://peagent.org/) integrating the full analytical framework described in this study (**Figure 5I**). For any user-submitted DNA sequence and a species model of the user’s choice, the portal returns cell-type-resolved chromatin accessibility predictions and per-nucleotide attribution maps, supports cell-type-level contribution pattern search, and enables JBrowse-based visualization of attribution scores across genomic loci. This lets users evaluate synthetic promoter or enhancer designs, or test how any mutation, motif insertion, disruption or deletion alters chromatin accessibility at cell-type resolution. Together, these functionalities position PEAgent as a unified resource for dissecting plant *cis*-regulatory sequence logic and interpreting noncoding genetic variation at cell-type level.

## Discussion

Plants have long lacked a systematic understanding of *cis*-regulatory grammar at cell-type resolution. PEAgent addresses this gap by predicting single-cell chromatin accessibility directly from DNA sequence and by rendering the learned regulatory logic interpretable at cell-type resolution, together across 320 cell types, at atlas scale and across soybean, maize and rice, providing a unified framework and data resource for training, evaluating and interpreting sequence-to-accessibility models in plants. Earlier plant sequence-based approaches have remained at the bulk level. PlantDeepSEA predicts chromatin accessibility from sequence but aggregates signal across whole tissues^39^. AgroNT, a plant genomic language model, has likewise been evaluated on tissue-level supervised tasks and does not preserve single-nucleotide resolution because it tokenizes sequences into 6-mers^18^. By training on such bulk measurements, these models average signal across heterogeneous cell populations, obscuring much of the cell-type-specific regulatory grammar. Their interpretation has also largely focused on individual high-impact sites, typically tissue-level base motifs, with the more complex combinatorial grammar of motif composition, orientation and spacing remaining less explored. PEAgent extends this line of work by resolving that combinatorial grammar at cell-type resolution.

We also developed an openly accessible, interactive PEAgent database (https://peagent.org) that rapidly predicts cell-type chromatin accessibility and attribution maps for user-defined sequences, making these models directly usable by the plant community. This is particularly valuable for experimental and synthetic biology. Users can submit any DNA sequence, select the best-matched species model, and obtain cell-type-resolved chromatin accessibility, enabling rapid *in silico* screening of synthetic promoter or enhancer designs and testing of how any mutation, motif insertion, disruption or deletion reshapes chromatin accessibility in specific cell types. To interpret these predictions, an *in silico* mutagenesis module returns global or cell-type-specific attribution maps that pinpoint the seqlets driving chromatin accessibility in a designed or edited sequence. Beyond this, an integrated JBrowse instance serves precomputed attribution maps for soybean, maize and rice and shows how motif combinations contribute differently to chromatin accessibility across cell types. We further provide a searchable module for the cell-type-resolved lexicon of all three species, reporting each pattern’s cell type of origin, contribution weight matrix, best-matched annotated motif, cross-species seqlet distribution and classification, together with a Python package (peagent-tool) for high-throughput prediction and attribution on large sequence sets. Together, these resources lower the barrier to sequence-based, cell-type-resolved regulatory analysis and make PEAgent a foundation on which the plant community can build.

We resolved a lexicon of 243 contribution patterns at cell-type resolution. Some of these recovered patterns were already apparent at the global level, with TCP and bHLH emerging as the strongest determinants of chromatin accessibility. This agrees with sequence-based models trained on bulk rice data^40^ and with maize caQTL in which variants at TCP motifs alter chromatin accessibility^24^, showing that PEAgent recovers established, global regulatory signals. These core patterns were also strongly conserved across the two monocots and the dicot examined. This conservation is consistent with cross-species DAP-seq analyses showing that transcription-factor binding preferences are deeply conserved across flowering plants^41^. More consequential, however, was the cell-type-resolved layer, which revealed many patterns not detected at the global level, nearly half of them composite motifs that combine two spatially distinct binding sites. A comparable predominance was reported in human development, where 64.8% of cell-type-resolved patterns were composite^15^, suggesting that combinatorial grammar is a general feature of cell-type-specific chromatin accessibility. A further strength of PEAgent is its ability to resolve this combinatorial grammar, whereas conventional motif-enrichment analyses identify individual motifs but cannot systematically assess how they combine in order, orientation and spacing.

Using co-occurrence and synergy analyses, we further resolved this grammar into two distinct modes of cooperative action. In the short-range mode (Type 1, ≤20 bp), adjacent transcription factors stabilize each other’s binding through direct contact or DNA-mediated allostery and cooperatively displace the local nucleosome^42–45^. Such short-range cooperativity has also been observed in animals. During human development cells, some composite motifs with fixed spacing and orientation reflect DNA-mediated cooperativity between directly interacting transcription factors^15^, and another sequence-footprint model identified analogous short-range composites, such as a Runx:Ets element whose direct interaction was supported by AlphaFold3^46^. The second mode (Type 2, 20–150 bp) acts within a single nucleosome and is consistent with collective transcription-factor competition against the nucleosome^28^. To our knowledge, such spacing-resolved cooperative grammar has not previously been systematically characterized in plants. By quantifying motif-pair synergy across different spacings and orientations at cell-type resolution, PEAgent provides a framework and resource for *in silico* testing these interactions.

The potential of sequence-to-function models remains incompletely explored. Most are used only to predict signals for a given sequence or to recover individual transcription-factor motifs, yet they are also powerful for studying regulatory evolution across species. Despite rapid sequence evolution, a deeply conserved regulatory code exists across plants, as recently mapped by alignment-based analysis of conserved noncoding sequences spanning ∼300 million years^8^. Such alignment-based approaches define where conserved sequences lie, but not whether a substitution preserves or disrupts regulatory activity. PEAgent adds this functional layer, predicting regulatory activity rather than scoring sequence similarity and identifying whether changes fall at positions critical for chromatin accessibility. Accordingly, PEAgent-predicted chromatin accessibility distinguished grass-conserved from rice-lineage-specific ACRs far more accurately than primary-sequence conservation scores (PhastCons and PhyloP) in this study, mirroring the advantage of sequence-based evolutionary models over conservation scores in mammalian CREs^14^. This parallels protein evolution field, where structure-prediction models recover conserved function among sequences too divergent for alignment^47^. Functional regulatory conservation is therefore not equivalent to primary-sequence conservation, and cell-type-resolved sequence models can trace regulatory histories that sequence alignment alone cannot resolve.

In summary, PEAgent brings sequence-based prediction of chromatin accessibility to cell-type resolution in plants. Beyond prediction, we dissect combinatorial regulatory grammar at the cell-type level and explore how predicted regulatory activity can trace cis-regulatory evolution across species. PEAgent gives the plant community a foundational resource for decoding cell-type-specific cis-regulatory logic.

## Limitations of the study

PEAgent currently comprises independently trained species-specific models for soybean, maize, and rice, each based on scATAC-seq atlas from a single growth condition. Extending their chromatin accessibility predictions or *cis*-regulatory grammar interpretation to other environmental conditions or more distant species warrants caution and additional training data. PEAgent currently predicts single-cell chromatin accessibility, but it lacks the capability to determine the function of individual enhancers or repressors in setting gene-expression levels. We have supported the model’s cell-type-resolved contribution patterns and cooperative grammar with footprint analysis and with the co-expression of the transcription factors underlying co-occurring motifs in independent scRNA-seq datasets, and we have validated its sequence variant-effect predictions against maize population scATAC-seq.

## Methods

### Plant single-cell ATAC-seq datasets and preprocessing

Model training and evaluation used and single-nucleus ATAC-seq (scATAC-seq) atlases that we previously generated for maize (*Zea mays*)^20^, rice (*Oryza sativa*)^21^, and soybean (*Glycine max*)^22^. The soybean atlas comprised 200,732 nuclei from 10 tissues (98 cell types, 303,199 ACRs), the maize atlas 50,632 nuclei from 6 tissues (94 cell types, 158,586 ACRs) and the rice atlas 104,029 nuclei from 9 organs (128 cell types, 128,764 ACRs). For each atlas, we used the 500-bp accessible chromatin region (ACR) definitions and cell-type annotations reported in the original studies, comprising 303,113 soybean, 158,586 maize, and 128,764 rice ACRs. ACR coordinates are defined from the corresponding species (soybean, Wm82.a4.v1; maize, B73 RefGen_v5; rice, Nipponbare MSU v7). For each species, we took the cell × ACR accessibility matrix and cell-type annotations from the atlas and binarized the matrix, so that each entry indicated whether an ACR was accessible in each cell. Each ACR was represented by a 1,344-bp sequence centred on its 500-bp peak, with positions extending beyond the chromosome padded with N, and one-hot encoded over the four nucleotides (A, C, G, T) as a 1,344 × 4 matrix. In each species, ACRs were randomly split into +training, validation, and test sets at a 90:5:5 ratio.

### Definition of cell-type-specific and constitutive ACRs

Cell-type-specific ACRs (ctACRs) for soybean and rice were obtained from the supplementary data of the published atlases^21,22^. For maize, which lacked a published ctACR set, we followed the approach used for soybean^22^ and defined ctACRs with SnapATAC2^48^ (marker_regions, P < 0.01), retaining peaks that were marker-called in no more than two cell types within each tissue.

Constitutive ACRs were defined from the remaining, non-cell-type-specific ACRs on the basis of how broadly they were accessible across pseudo-bulk groups, using a Bayesian-smoothed accessibility estimate to regularize groups with few cells. For each ACR in each group (defined by tissue and cell type, with replicate where available), accessibility was estimated as a Beta-binomial posterior mean, p^ = (k + α)/(n + α + β), where n is the number of cells in the group, k the number with non-zero accessibility and α = β = 1 (a uniform prior). An ACR was called open in a group when p^ exceeded 0.05, and its open ratio was defined as the fraction of groups in which it was open. ACRs whose open ratio exceeded a species-specific cutoff (0.80 in soybean, 0.85 in rice and 0.60 in maize) were classified as constitutive. We calibrated the cutoff separately for each species, because differences in sequencing depth and cell-type sampling shifted the open-ratio distribution and precluded a single cutoff from yielding comparable constitutive sets.

### Model architecture and training

PEAgent is a sequence-based convolutional neural network adapted from scBasset that predicts single-cell chromatin accessibility directly from DNA sequence^12^, mapping a one-hot-encoded sequence through a convolutional tower to a dense bottleneck and a final sigmoid output layer. We retained the published architecture with a bottleneck of 64 units and an output layer sized to the number of cells profiled in each species, so that the model predicts, for every input sequence, a vector of per-cell accessibility probabilities (ŷ ∈ [0,1]^n^ ^cells^) as a multi-task classification across all single cells.

A separate model was trained for each species. Training sequences were augmented by stochastic reverse-complementation and random shifts of up to 3 bp, and model parameters were optimized against a binary cross-entropy loss with the Adam optimizer (learning rate 0.01, β₁ = 0.95, β₂ = 0.9995) and a batch size of 256 for up to 100 epochs. Training was monitored by the multi-label area under the receiver operating characteristic curve (AUROC) and area under the precision-recall curve (AUPR); the best checkpoint was retained by maximum training AUROC, and early stopping was applied (patience 50, minimum improvement 1 × 10⁻⁶) with restoration of the best weights.

### Model evaluation

Model performance was evaluated from four perspectives: held-out ACR prediction accuracy, Correction between predicted and observed cell type chromatin accessibility profiles, performance across ACR classes, and the biological information of model-learned cell embeddings.

1. For held-out prediction accuracy, models were evaluated on ACRs that were not used during training from the testing set. Held-out prediction accuracy was assessed using AUROC and AUPR at both the cell-group level, across held-out ACRs, and per-ACR level. As a negative control, peak labels were shuffled across held-out ACRs while input sequences were kept unchanged.
2. Quantitative agreement between predicted and observed accessibility was assessed by computing PCCs between cell-type-specific pseudo-bulk prediction profiles and observed accessibility profiles across held-out ACRs.
3. Context-dependent performance was assessed by comparing AUROC between cell-type-specific and constitutively accessible ACRs, as well as across genomic contexts, including genic, proximal, and distal ACRs. Genomic categories were assigned from the position of the ACR summit relative to the nearest annotated gene, as genic (within a gene body), proximal (≤2 kb) or distal (>2 kb).
4. We assessed the biological content of these model-learned embeddings, defined as the per-cell weights of the final dense layer, both qualitatively and quantitatively. Qualitatively, UMAP visualization revealed whether cells organize by tissue and cell type in the learned embedding space. Quantitatively, we grouped cells by their embeddings using Leiden clustering and measured how closely these unsupervised, sequence-derived clusters matched the annotated cell identities with normalized mutual information (NMI)^49^, a score ranging from 0 (no correspondence) to 1 (perfect correspondence).

### Contribution pattern analysis

For each 1,344-bp ACR window, we performed *in silico* mutagenesis at the model bottleneck layer by substituting each position with all four nucleotides and recording the resulting change in the 64-dimensional bottleneck activation. This produced a position-by-nucleotide-by-bottleneck-dimension mutation-effect tensor for each ACR.

#### Bulk-level and cell-type-level attribution maps

To convert bottleneck-level effects into nucleotide-resolution attribution maps, bottleneck ISM tensors were projected through the final dense layer. For each target group *g*, where *g* denotes either all cells for global attribution or a specific tissue–cell-type group, the corresponding output-layer weight vector was defined as the average final-layer weight across cells in that group:

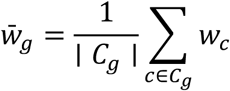

The mutation score for each position *i* and substituted nucleotide *b* was then computed by projecting the bottleneck mutation effect onto this group-level weight vector:

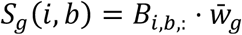

Thus, global attribution maps represent mutation effects averaged across all cells, whereas cell-type-level maps represent effects averaged within each tissue–cell-type group.

Then hypothetical contribution and reference-base attribution maps were generated at both global and cell-type levels for downstream contribution pattern discovery. For each projected mutation score matrix, hypothetical contribution scores were obtained by centering the A/C/G/T scores at each position:

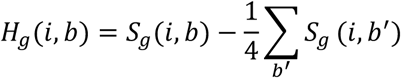

Hypothetical contribution maps therefore retained the centered effects of all four possible nucleotides, whereas reference-base attribution maps retained only the contribution of the observed genomic base, with all non-reference bases set to zero. Global tracks were used after per-position centering. For cell-type-specific tracks, tissue-level background was additionally corrected by subtracting the mean hypothetical contribution across all valid cell types within the same tissue:

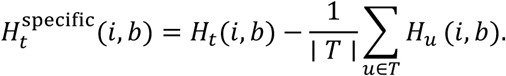

These tissue-normalized cell-type tracks were used for cell-type-resolved contribution pattern discovery.

#### Global contribution patterns and genome-wide instances

To identify global contribution patterns, TF-MoDISco^23^ was applied to global level attribution maps from constitutively accessible ACRs. This recovered dominant sequence patterns for each species as contribution-weight matrices (CWMs). These CWMs were then mapped back to all ACR attribution tracks using Fi-NeMo (https://github.com/kundajelab/Fi-NeMo), which competitively assigned motif instances under an L1 penalty. Genome-wide instances were called across all ACRs in soybean, maize, and rice, and unique instance counts were summarized for each pattern at (λ=0.7).

#### Cell-type-level contribution patterns

To identify cell-type-level contribution patterns, TF-MoDISco was run independently on cell-type-specific attribution maps from ctACRs for each cell type, yielding 2,120 raw contribution patterns. To reduce redundancy across cell types and species, raw CWMs were clustered into meta-patterns using MotifCompendium (https://github.com/kundajelab/MotifCompendium) based on CWM similarity. Clustering was performed using Leiden clustering with the CPM objective, a similarity threshold of 0.9 and a resolution of 1.0. Only contribution patterns with positive attribution signals were retained for downstream analysis. After removal of noisy CWMs characterized by diffuse contribution, GC-rich signal, or lack of dominant base preference, final curation yielded 243 contribution meta patterns at cell-type level.

#### Motif annotation

Each resulting meta-pattern CWM was annotated with TomTom^50^ using a plant motif database containing 1,453 motifs, including 927 non-redundant JASPAR 2026 CORE plant motifs^51^ and 526 Arabidopsis DAP-seq motifs^4^. TomTom was run with Pearson distance and a minimum overlap of 5 bp, and the best centered hit with the lowest q value was retained for each trimmed CWM core.

#### Classification and labelling of cell-type level mata-pattern

Each meta-pattern was classified using CWM-derived features, including the motif core trimmed at 30% of the maximum per-position L1 norm, the number and position of L1 sub-peaks, and the flank ratio, defined as the fraction of core L1 signal outside a ±8-bp primary-peak window. Base patterns were defined as single-site motifs with low flanking contribution (flank ratio ≤0.15) and a short trimmed core (≤13 bp). Base with flank patterns contained one dominant motif site with broader surrounding contribution. Homocomposite and heterocomposite patterns contained two spatially distinct *cis*-regulatory zones corresponding to the same or different TF families, respectively. Unresolved patterns showed concentrated sequence signal but lacked a confident motif match (TomTom q ≥ 0.10). All meta-patterns were manually reviewed. Single-site patterns were labelled as <id>.<TF family>:<TF-best-hit># <INDEX>. Composite patterns were labelled as <id>.<FAMILY1>:<TF1>_<FAMILY2>:<TF2># <INDEX>. Unresolved patterns were labelled as <id>.unresolved. <FAMILY>-like# <INDEX>. For composite patterns, labels were assigned according to the two spatially distinct sub-motifs ordered from the 5′ zone to the 3′ zone. Pattern indices were ordered by total seqlet support, and all labels and zone assignments were manually verified.

### Footprint analysis

Motif-centered footprint analysis was performed for the unresolved bZIP-like P5 metapattern. Maize leaf scATAC-seq reads were aligned with Chromap^52^, with Tn5 cut-site shift correction applied during alignment. Shift-corrected fragments were aggregated into pseudo-bulk profiles and imported into scPrinter^46^. Sequence-dependent Tn5 bias was corrected using the scPrinter bias model, from which we generated a maize genome-wide predicted bias track based on the B73 v5 reference genome. Multiscale footprint scores were computed for each seqlet in an 800-bp motif-centered window across scales 4–50. For aggregate visualization, profiles were oriented to the same motif strand, averaged across anchors, and displayed within an approximately 100-bp motif-centered region.

### Contribution pattern co-occurrence analysis

For each cell-type-resolved base meta-pattern, source seqlets were retrieved from the TF-MoDISco HDF5 output, including their originating ctACR, genomic position, and cell-type annotation. Co-occurrence was assessed for pairwise combinations of the 57 base meta-patterns by identifying cases in which two seqlets occurred within the same ACR. For each co-occurring pair instance, motif spacing was recorded as the center-to-center distance between the two seqlets, which reduces bias caused by differences in motif length. Across all cell types, this analysis examined 1,596 pairwise combinations and identified 261,366 co-occurrence instances. For each instance, we recorded the cell type of the corresponding ctACR, the anatomical category of that cell type, whether the pair was homotypic or heterotypic, and the center-to-center distance. For each meta-pattern pair, the median center-to-center distance and deviation were summarized across all co-occurrence instances.

### Synergy analysis

Synergy was quantified using an adapted *in silico* marginalization strategy^15^ and the corresponding species-specific PEAgent model. To ensure adequate statistical power and biological relevance, we tested only base-motif pairs with at least 20 co-occurrence instances in their most abundant cell type, and each pair was evaluated in that cell type, yielding 167 pairs, including 143 heterotypic and 24 homotypic pairs. For a cell-type group *g*, model prediction was read out as a pseudo-bulk accessibility logit,

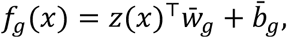

where *z*(*x*) is the bottleneck embedding of sequence 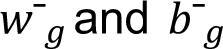 are the final-layer output weights and bias averaged across cells in group *g*. Thus, *f*_g_(*x*)represents the predicted pre-sigmoid accessibility of sequence *x* in that cell type. For each species, 100 non-accessible 1,344-bp background sequences were sampled from non-peak regions and GC-matched to ACR windows.

Motif effects were defined as changes in predicted accessibility after motif insertion into the center of each background sequence. For motifs *A* and *B*, the single-motif effects, joint effect, and additive expectation were defined as 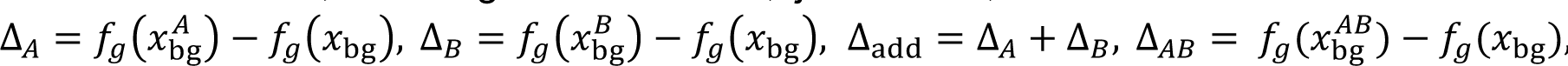 The synergy score was then defined as

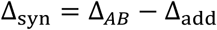

Joint effects were evaluated across motif orientations and spacings. Heterotypic pairs were tested in four relative orientations (*A*^+^*B*^+,^ *A*^+^*B*^−,^ *A*^−^*B*^+,^ *A*^−^*B*^−^), whereas homotypic pairs were tested in three non-redundant orientations. edge-to-edge spacing was scanned from 0-200bp. The center-to-center (c2c) and edge-to-edge (e2e) distances between motif pairs were related as follows: 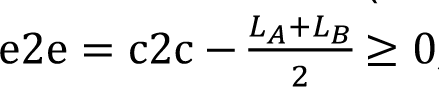, where *L_A_*and *L_B_* are the motif lengths. For each eligible arrangement, Δ_syn_was computed across the 100 paired background sequences. For each arrangement of each motif pair, we tested whether Δ_syn_ > 0 using a one-sided Wilcoxon signed-rank test across the 100 backgrounds, followed by a single Benjamini–Hochberg correction across all tested arrangements from all 167 pairs. Each pair was summarized by its optimal arrangement, defined as the eligible arrangement with the largest mean joint effect Δ*_AB_*. The adjusted *P* value and mean Δ_syn_ of this arrangement were used as the pair-level significance and synergy magnitude, respectively. Pairs with adjusted *P* < 0.01 were called synergistic, yielding 59 synergistic pairs, and were classified by their optimal edge-to-edge gap as Type 1 (<20 bp), Type 2 (20–150 bp), or distal (>150 bp). To confirm that predicted synergy was driven by the inserted motifs rather than flanking sequence, attribution maps were computed in the tested cell type for both the background sequence and the optimal composite sequence. Attribution was examined over the window spanning the two motifs, and synergistic predictions were retained only when accessibility-predictive positions coincided with the inserted motifs.

### Cross-species prediction and evaluation

We tested whether PEAgent models transfer across species by predicting ACR accessibility among soybean, maize and rice at both bulk and cell-type resolution. Nine directed source-to-target combinations were evaluated, comprising the six cross-species directions among the three species and three within-species baselines. For each cross-species combination, source-species models were evaluated on target-species ACR sequences by auROC, using observed target-species chromatin accessibility as ground truth. For cell-type-level comparisons, only tissue–cell-type pairs with matching annotations in both source and target species were included.

As an independent test of cross-species generalization, we used the rice PEAgent model to predict accessibility for the ACR sequences of a maize crown-root and leaf scATAC-seq atlas. The predicted and observed cell-by-ACR matrices were each processed with the standard SnapATAC2 pipeline for dimensionality reduction, clustering and visualization. We then compared the predicted and observed data in two ways, by testing whether their unsupervised clustering recovered known maize cell identities on UMAP, and by comparing the deviations of key transcription-factor motifs between the two.

### Grass-conserved and lineage-specific ACR prediction

Rice ACRs were classified by cross-species conservation status using the published grass leaf scATAC atlas^21^, which reports synteny-based ACR comparisons across rice and four other grasses: *Zea mays*, *Sorghum bicolor*, *Panicum miliaceum* and *Urochloa fusca*. For each rice ACR, the atlas annotated whether syntenic regions were present in the other four grass genomes and whether those regions were accessible. Rice ACRs were labelled as grass-conserved when all four syntenic regions in the other grasses were called as ACRs, and as rice-specific when all four syntenic regions were present but not accessible. Grass-conserved ACRs were assigned label 1, and rice-specific ACRs were assigned label 0.

For PEAgent-based prediction, each rice ACR and its corresponding syntenic regions in the four other grasses were treated as one syntenic sequence group. For each sequence in the group, a 1,344-bp window centered on the syntenic region was extracted from the corresponding species genome and scored using the rice PEAgent model. Only outputs from leaf-related cell types were used. Prediction scores were averaged across available sequences from 5 grass species within each syntenic group to generate an ACR-level PEAgent score. Two PEAgent summary scores were calculated: the mean prediction across all leaf cell types and the maximum prediction across leaf cell-type groups.

As conservation baselines, we used precomputed genome-wide phastCons scores from PlantRegMap^36^ and grass-level phyloP^37^ scores projected onto rice genomic coordinates. For each selected rice ACR, mean phastCons and phyloP scores were calculated across the rice ACR interval from the corresponding bedGraph conservation tracks. Prediction performance was evaluated by auROC for each continuous score against the binary grass-conserved versus rice-specific labels. Confidence intervals were estimated by 1,000 stratified bootstrap resamplings of positive and negative ACRs, preserving class balance in each resampling to assess the robustness of classification performance.

### Variant effect prediction and validation with caQTL data

We adapted the AlphaGenome^11^ chromatin-accessibility variant-effect validation strategy to fine-mapped maize caQTLs at cell-type resolution^24^. In total, 46,443 fine-mapped *cis*-caQTL SNVs from 172 diverse maize inbred lines were analyzed across 13 seedling cell types that could be matched to PEAgent outputs, with each SNV annotated by posterior inclusion probability (PIP), caQTL effect size, and REF/ALT alleles. For each SNV located within an ACR, REF and ALT input sequences were generated with the variant centered in the model input window and differing only at the variant position. Both alleles were scored with the maize PEAgent model using the corresponding matched cell-type output, and predicted variant effects were calculated as the signed change in pre-sigmoid accessibility logit, Δlogit = *f*_g_(ALT) − *f*_g_(REF). We compared two model settings: the zero-shot maize PEAgent model and a B73 seedling model fine-tuned from the maize atlas model using seedling cell-type accessibility labels. Fine-mapped caQTL SNVs with PIP ≥ 0.1 were treated as candidate causal variants. For each candidate causal SNV, four matched non-causal control SNVs were selected from the nominal *cis*-caQTL variant set, requiring the same ACR and cell type, valid REF/ALT alleles, and absence from the fine-mapped causal set in that cell type. For each cell type, we tested whether candidate causal SNVs showed larger predicted effect magnitudes than matched same-ACR controls, using ∣ Δlogit ∣as the effect score. Candidate effects were compared with the mean effect of their four matched controls using a one-sided Wilcoxon signed-rank test, followed by Benjamini–Hochberg correction across cell types.

## Data and code availability

All scATAC-seq, scRNA-seq raw datasets analyzed in this study were obtained from publicly available repositories. Soybean scATAC-seq and scRNA-seq data were obtained from GSE270392 and PRJNA1124403^22^; rice scATAC-seq data from PRJNA1007577/GSE252040^21^; maize scATAC-seq data from GSE155178^20^ and population-scale scATAC-seq data from 172 genetically and phenotypically diverse maize inbred lines from GSE275410^24^; and comparative grass scATAC-seq data from PRJNA1063172^37^. PEAgent models and attribution maps generated in this study are available through the PEAgent web portal (https://peagent.org). Any additional information required to reanalyze the data reported here is available from the lead contact upon request. All code used to train, evaluate and interpret PEAgent models, together with the peagent_tools Python package for sequence prediction and attribution analysis, is publicly available at https://github.com/YAOJ-bioin/peagent_tool.

## Acknowledgements

This material is based upon work supported by the U.S. Department of Energy, Office of Science, Biological and Environmental Research Program under award number DE-SC0025995, by the National Science Foundation (IOS-2134912 and DBI-2516239) and the UGA Office of Research to RJS. We also thank the Georgia Advanced Computing Resource Center (GACRC) at the University of Georgia for computational resources.

## Author contributions

J.Y. led the study and contributed to all major components, including framework design, data analysis, database construction, visualization, and writing of the original manuscript. R.J.S. guided the overall direction of the study, contributed to analytical discussions, manuscript writing and revision, and supervised the project. J.L. contributed to analytical discussions and manuscript revision. X.Z. and X.L. contributed to discussions of specific analyses. E.P. and A.P.M. contributed to project discussion and manuscript revision. All authors reviewed and approved the final manuscript.

## Supplemental information

**Figure S1.**
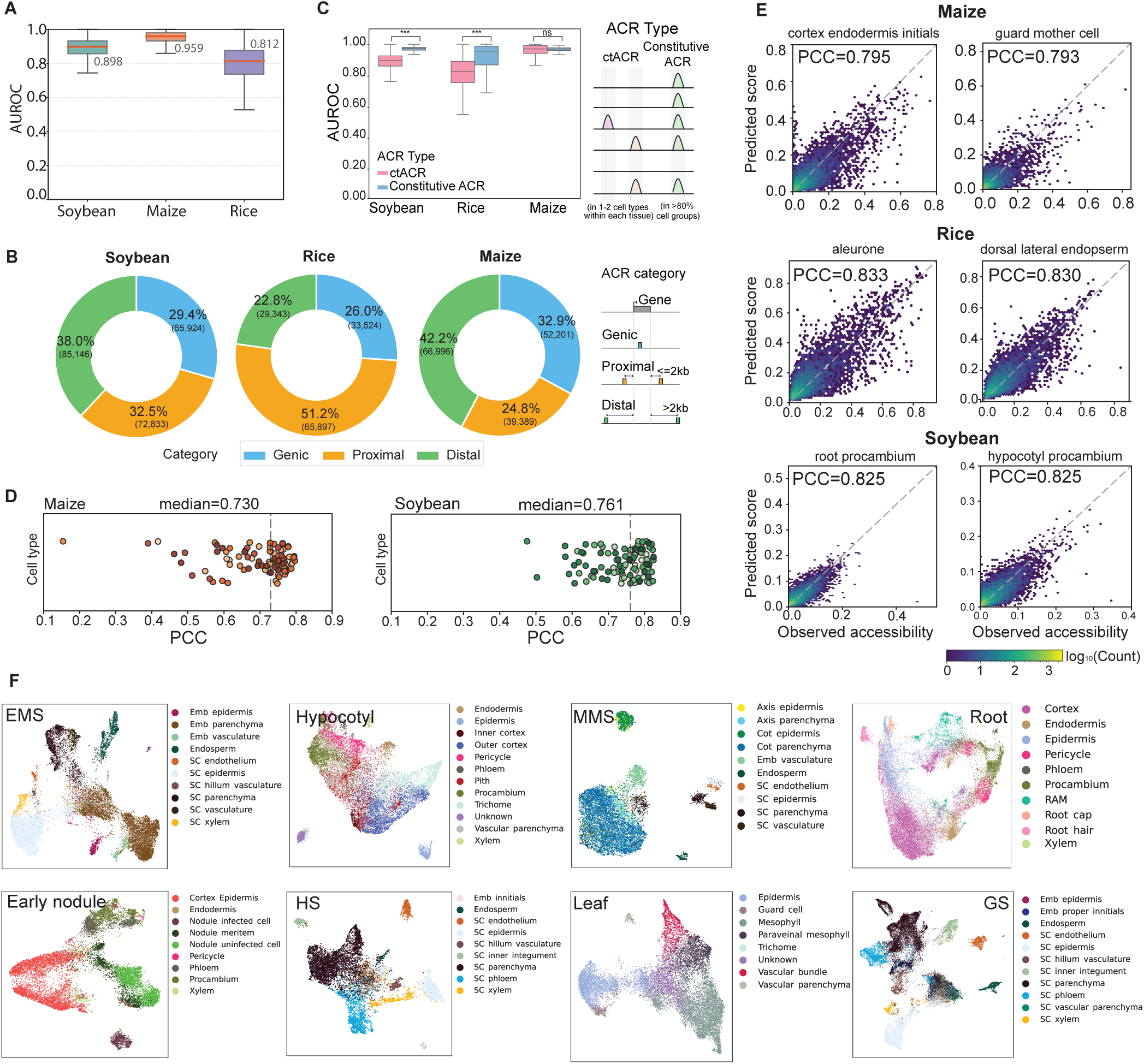
Extended evaluation of PEAgent performance, related to Figure 1. **A,** Per-peak AUROC on held-out ACRs across species. **B,** Proportions of genic, proximal, and distal ACRs in each species. **C,** AUROC for cell-type-specific (ctACRs; accessible in 1–2 cell types per tissue) and constitutive A CRs (>80% of cell groups) across species. Wilcoxon rank-sum test; ns, not significant; ***P < 0.001. **D**, Per-cell-type PCC between predicted and observed chromatin accessibility on held-out peaks in maize and soybean. Each dot represents a cell type **E**, Representative scatter plots of predicted versus observed accessibility on held-out peaks for individual cell types across species. Color indicates point density. **F,** UMAP of model-learned cell embeddings for individual soybean tissues, colored by annotated cell type. EMS, early maturation stage; HS, heart stage; MMS, middle maturation stage; GS, globular stage.

**Figure S2.**
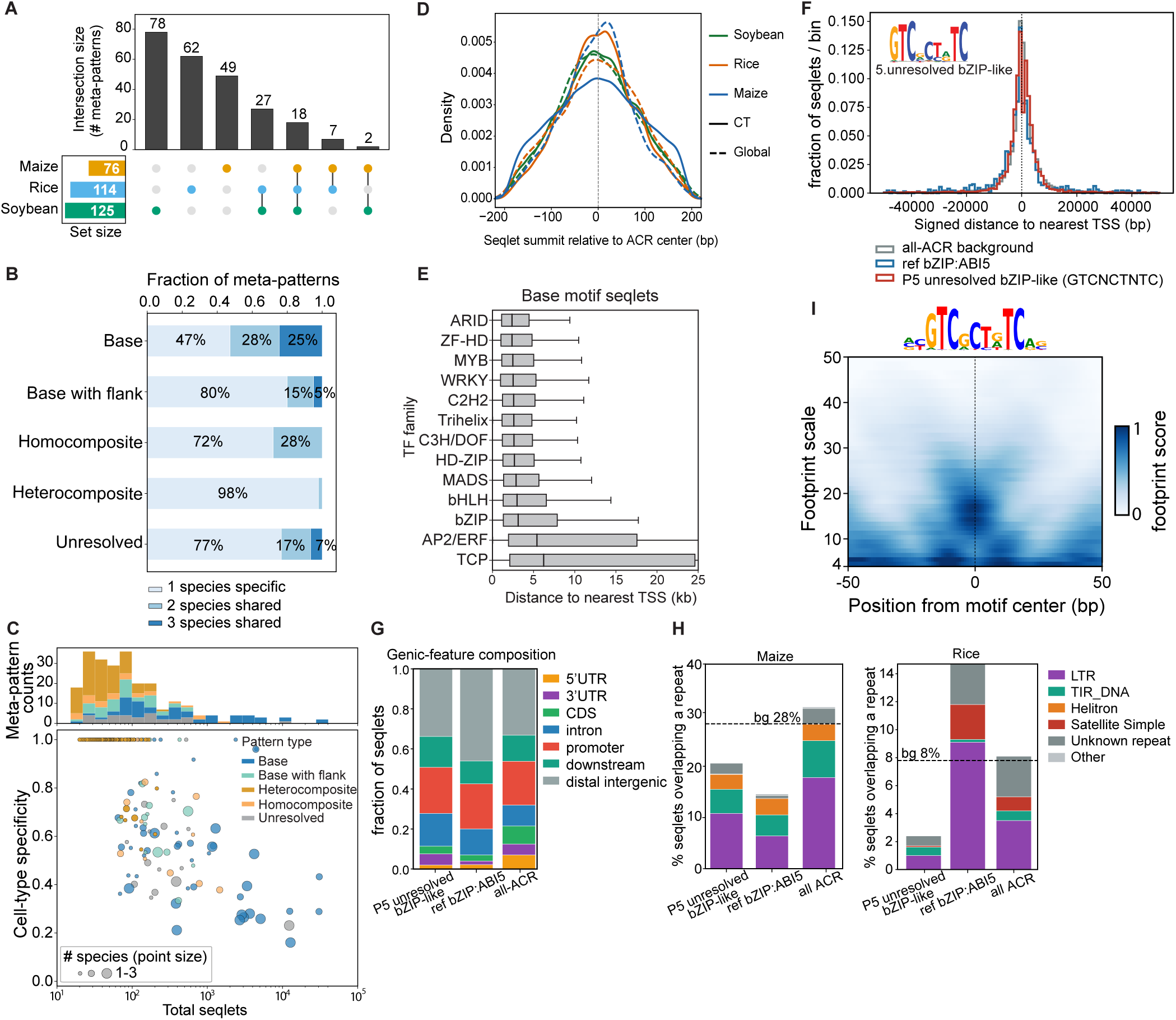
Conservation, genomic distribution and unresolved pattern properties of PEAgent contribution meta-patterns, related to Figure 2. **A**, UpSet plot showing the species distribution and overlap of cell-type-resolved meta-patterns across maize, rice and soybean. **B,** Fraction of meta-patterns from each category that are species-specific or shared by two or three species. **C,** Relationship between total seqlet support and cell-type specificity for cell-type-resolved meta-patterns. Top, meta-pattern count distribution; bottom, each point represents one meta-pattern, coloured by pattern class and sized by the number of species in which it was detected. **D,** Density of seqlet summit positions relative to ACR centres for cell-type-resolved and global contribution patterns in each species. **E,** Distance from seqlets of cell-type-resolved base meta-patterns to the nearest transcription start site (TSS), grouped by TF family. **F-H,** Genomic annotation of P5 unresolved pattern seqlets (GTCNCTNTC) compared with reference bZIP:ABI5 seqlets and the all-ACR background, including signed distance to the nearest TSS (**F**), genic-feature composition (**G**) and repeat-class overlap in maize and rice (**H**); dashed lines indicate the all-ACR background. **I,** Multiscale footprint aggregated across seqlets instances of the unresolved pattern (consensus GTCNCTNTC) in maize leaf, after correction for Tn5 sequence bias.

**Figure S3.**
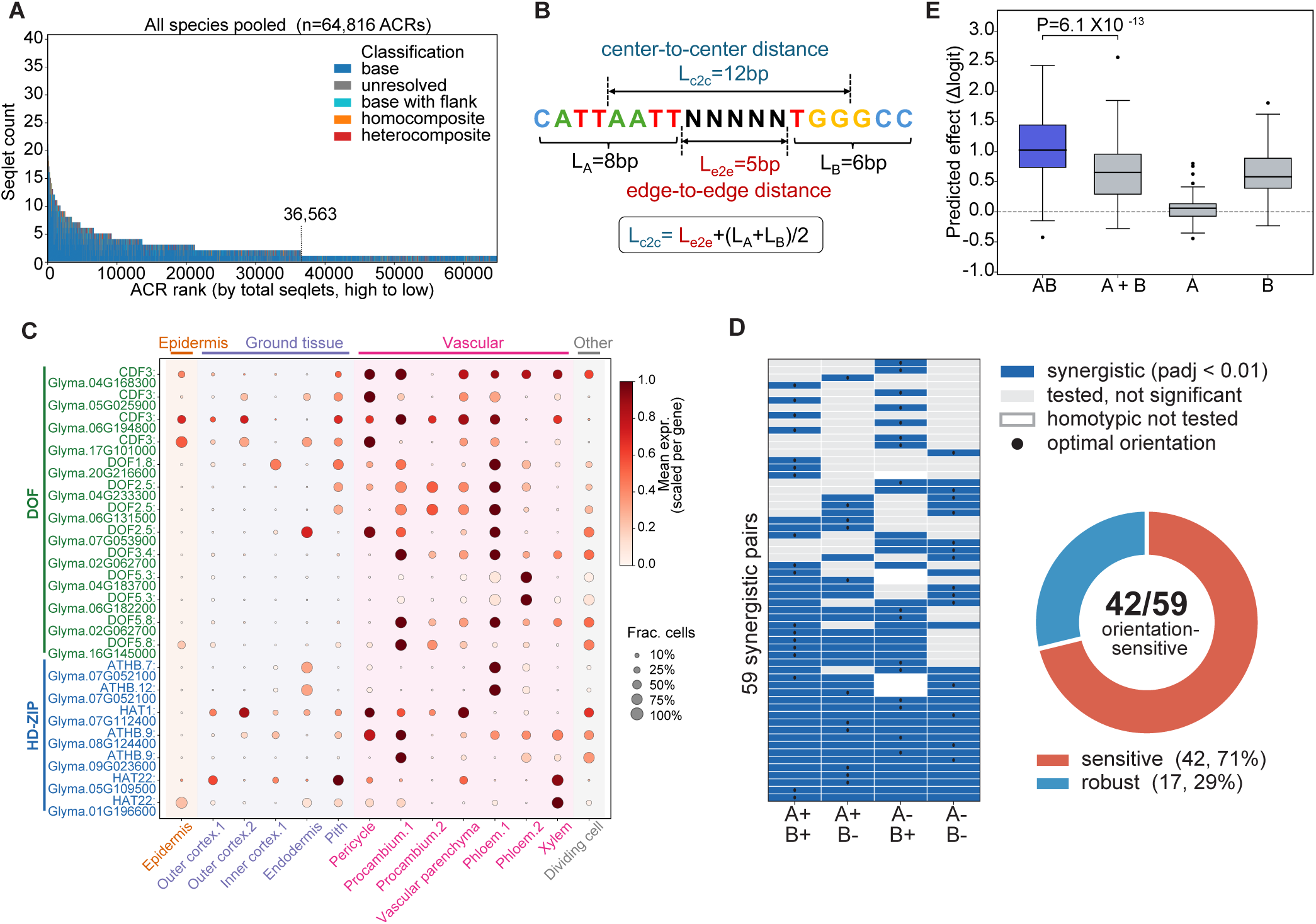
Co-occurrence prevalence, distance metrics and synergy characterization, related to Figure 3. **A**, Distribution of seqlet counts per ACR, ranked from high to low; bars are coloured by pattern classification (base, base-with-flank, homocomposite, heterocomposite and unresolved). **B**, Definitions of the centre-to-centre (c2c) and edge-to-edge (e2e) distances between two motifs and their conversion. **C**, Single-cell RNA-seq expression of representative DOF and HD-ZIP family TFs across cell types of the soybean hypocotyl. **D**, Orientation sensitivity of the 59 synergistic pairs. Heatmap shows significance across the four relative orientations (blue, synergistic at q value < 0.01). **E**, Predicted effects (Δlogit) of the HD-ZIP and TCP pair at its optimal arrangement, computed across 100 background sequences. AB, both motifs inserted jointly. A+B, additive expectation (sum of the individual effects). A and B, each motif inserted alone.

**Figure S4.**
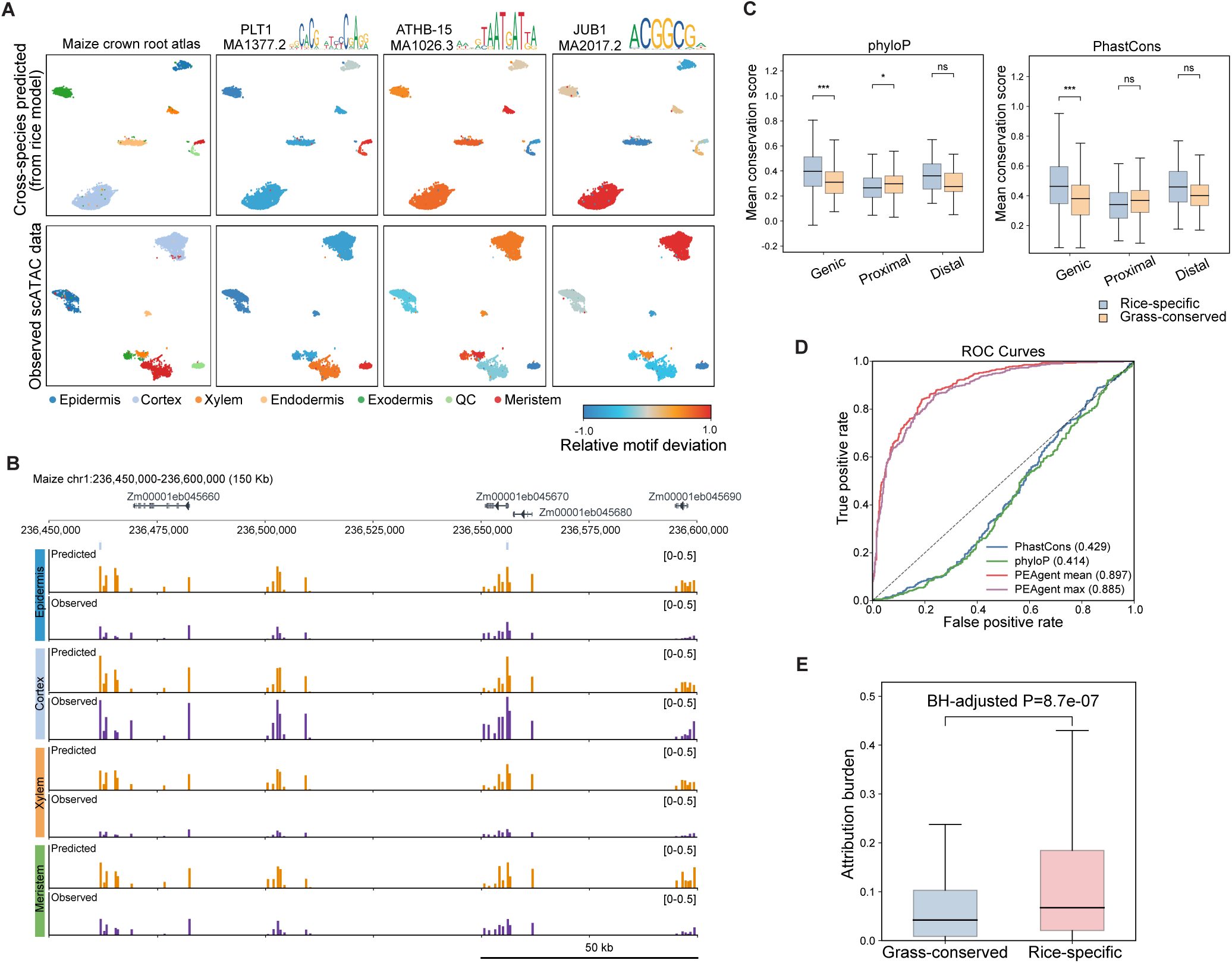
Cross-species chromatin accessibility prediction and evolutionary classification of rice ACRs, related to Figure 4. **A**, UMAP embeddings of predicted (top) and observed (bottom) maize crown root chromatin accessibility, colored by cell type or relative TF motif deviation scores. **B**, Genome browser tracks comparing predicted (rice-to-maize) and observed cell-type-resolved chromatin accessibility at a representative 150-kb locus on maize chr1 (236,450,000–236,600,000). Predictions were generated only at annotated ACR windows (each 500 bp), not continuously along the locus; each predicted signal therefore represents one 500-bp ACR scored by the rice PEAgent model. **C**, Mean phyloP (left) and PhastCons (right) conservation scores for rice-specific and grass-conserved ACRs stratified by genomic context (Genic, Proximal, Distal). **D**, ROC curves for classifying rice-specific versus grass-conserved ACRs using sequence conservation scores (PhastCons, phyloP) or PEAgent prediction scores (mean and maximum across leaf cell types in grass syntenic sequences). AUROC values in parentheses.

**Figure S5.**
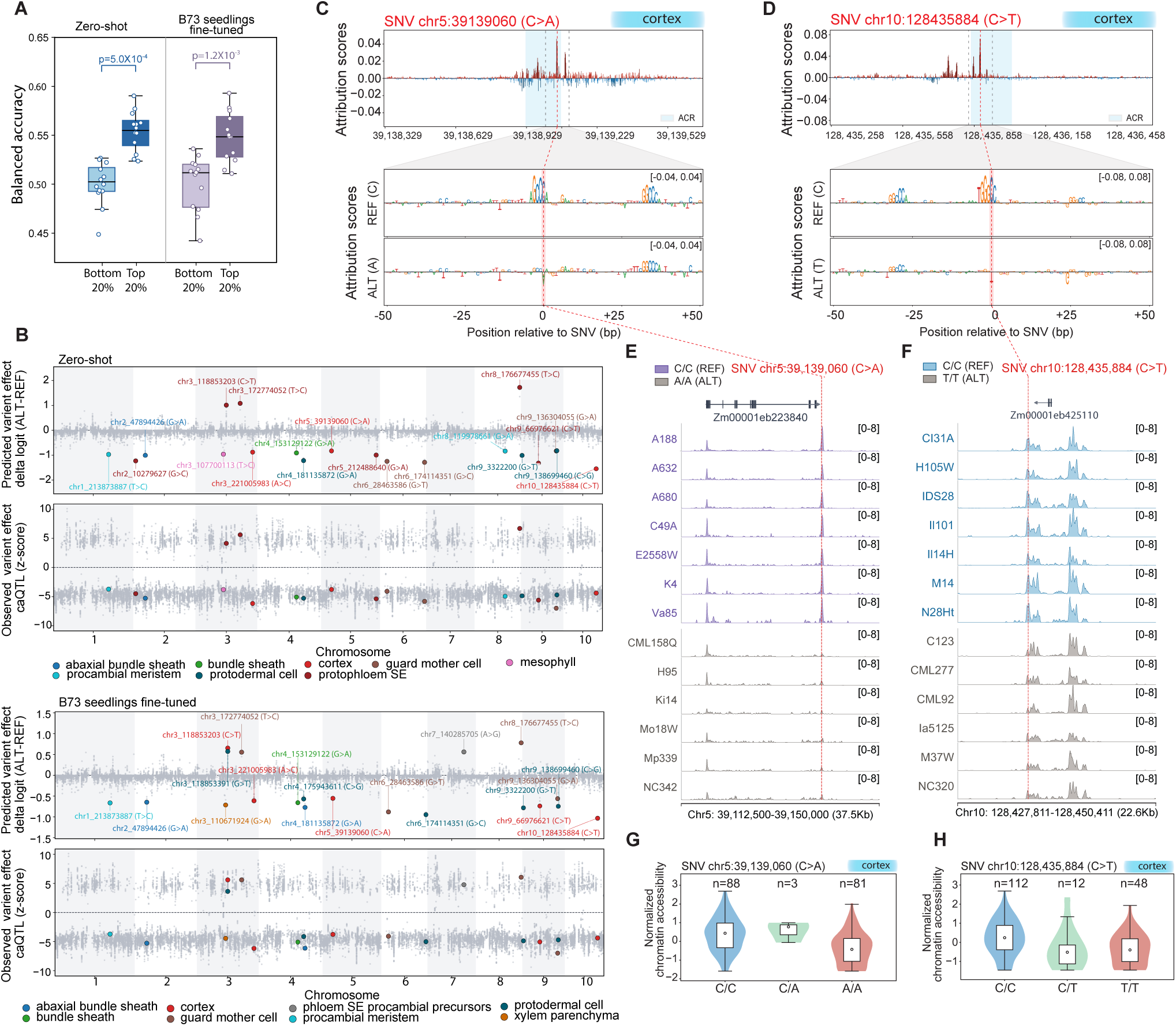
Validation of PEAgent variant effect predictions using maize population-level single-cell chromatin accessibility data, related to Figure 5. **A**, Balanced accuracy of predicted caQTL effect direction across cell types for the top 20% and bottom 20% fine-mapped caQTL SNV sites, using the zero-shot maize model (left) and the B73 seedling fine-tuned model (right). **B**, Genome-wide view of PEAgent-predicted variant effect and observed fine-mapped caQTL effects across cell types. Colored points highlight representative top 20 fine-mapped caQTL SNVs associated with individual cell types. Results are shown for both the zero-shot (top panel) and B73 seedling fine-tuned models (bottom panel). **C-D**, Attribution score profile for a cortex caQTL (SNV chr5:39,139,060, C>A) and (SNV chr10:128,435,884, C>T). Top: attribution scores across the 1,344-bp ISM scanning window, with the population-defined ACR highlighted in blue. Bottom: per-nucleotide attribution scores for REF and ALT alleles within a ±50-bp window, showing the substitution disrupts a binding motif at a high-attribution position. **E-F**, Genome browser tracks of chromatin accessibility at the chr5:39,139,060 and chr10:128,435,884 loci for representative maize genotypes grouped by REF and ALT alleles. Tn5 integrations are scaled per million. **G-H**, Normalized chromatin accessibility stratified by genotype at SNV chr5:39,139,060 and SNV chr10:128,435,884 across maize inbred lines.

**Table S1.** Cell-type-level prediction performance of PEAgent, including held-out AUROC and predicted–observed chromatin accessibility correlation, related to Figure 1.

**Table S2.** Cell-type-resolved contribution meta-patterns comprising the plant regulatory lexicon across soybean, maize and rice, related to Figure 2.

**Table S3.** *In silico* synergy analysis of co-occurring motif pairs, each tested in its most abundant cell type, related to Figure 3.

**Table S4.** Synergy analysis of the co-occurring motif pairs tested in endosperm, related to Figure 3.

**Table S5.** Cross-species conservation classification of rice ACRs with PEAgent prediction and sequence-conservation (PhyloP and PhastCons) scores, related to Figure 4.

